# Quantifying protein unfolding kinetics with a high-throughput microfluidic platform

**DOI:** 10.1101/2025.01.15.633299

**Authors:** B. Atsavapranee, F. Sunden, D. Herschlag, P.M. Fordyce

**Author notes:** These authors contributed equally to this work.

## Abstract

Even after folding, proteins transiently sample unfolded or partially unfolded intermediates, and these species are often at risk of irreversible alteration (*e.g.* via proteolysis, aggregation, or post-translational modification). Kinetic stability, in addition to thermodynamic stability, can directly impact protein lifetime, abundance, and the formation of alternative, sometimes disruptive states. However, we have very few measurements of protein unfolding rates or how mutations alter these rates, largely due to technical challenges associated with their measurement. To address this need, we developed SPARKfold (Simultaneous Proteolysis Assay Revealing Kinetics of Folding), a microfluidic platform to express, purify, and measure unfolding rate constants for >1000 protein variants in parallel via on-chip native proteolysis. To demonstrate the power and potential of SPARKfold, we determined unfolding rate constants for 1,104 protein samples in parallel. We built a library of 31 dihydrofolate reductase (DHFR) orthologs with up to 78 chamber replicates per variant to provide the statistical power required to evaluate the system’s ability to resolve subtle effects. SPARKfold rate constants for 5 constructs agreed with those obtained using traditional techniques across a 150-fold range, validating the accuracy of the technique. Comparisons of mutant kinetic effects via SPARKfold with previously published measurements impacts on folding thermodynamics provided information about the folding transition state and pathways via φ analysis. Overall, SPARKfold enables rapid characterization of protein variants to dissect the nature of the unfolding transition state. In future work, SPARKfold can reveal mutations that drive misfolding and aggregation and enable rational design of kinetically hyperstable variants for industrial use in harsh environments.

## Introduction

Most proteins must fold to function, so that thermodynamic stability—favoring the folded state—is required. But even when this criterion is met, proteins can still repeatedly unfold and sample higher free-energy conformations before refolding again^1,2^, and each excursion to an unfolded state proteins risks a variety of potential irreversible alterations, including cleavage by proteases, misfolding into an inactive conformation, and aggregation^3–8^. Some proteins prone to aggregation (*e.g.* transthyretin^9^) have evolved to be kinetically hyperstable with very slow unfolding rates, reducing the number of times they risk degradation or aggregation over cellular timescales^10^. Many secreted proteins, in addition to requiring stability to function in harsh environments, have similarly evolved kinetic stability to avoid irreversible unfolding.^10,11^ For proteins with modest kinetic stabilities, time-dependent degradation can serve as a molecular timer (*e.g.* serpin protease inhibitors and heat shock transcription factors)^12–17^. In these cases, post-translational modifications often alter unfolding rates; e.g., ubiquitination can increase unfolding rates to drive proteosomal degradation of target proteins^18,19^ while phosphorylation and methylation can modulate kinetic stability by altering protein interactions and conformational flexibility^20,21^. In an energy landscape diagram, thermodynamic stability is visualized as the difference in free energy between the folded and unfolded states and kinetic stability is visualized as the height of the transition state barrier between them (**Figure 1A**); under conditions in which refolding is slow compared to rates of cleavage or aggregation, changes in kinetic stability alone can drive significant changes in protein abundance over time^11^ (**Figure 1B**).

**Figure 1.**
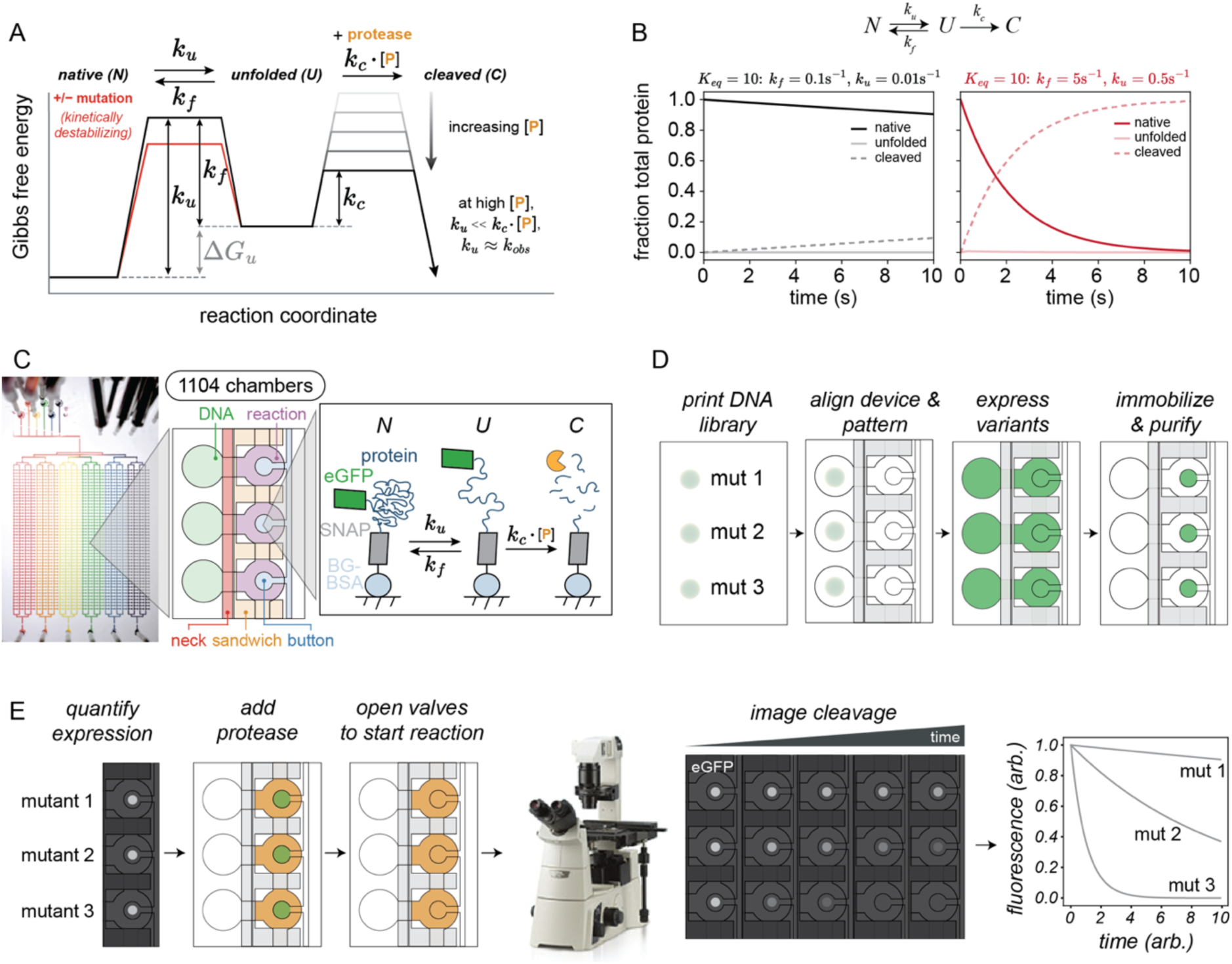
Protein folding and SPARKfold assay overview. **(A)** Gibbs free energy vs. reaction coordinate diagram for native, unfolded, and cleaved protein populations. Free energy of unfolding (Δ*G_u_*) is the difference in Gibbs free energy for the native and unfolded states (gray). Height of the energy barrier between the native and unfolded states depends on the unfolding rate (*k_u_*) and folding rate (*k_f_*), which is altered by kinetically destabilizing mutations (red); height of the energy barrier between the unfolded and cleaved states is defined by the cleavage rate constant (*k_c_*) and protease concentration ([P]). At high [P], *k*_u_ can be determined from the observed rate of proteolysis (*k_obs_*), as carried out herein. At low [P], *K*eq can be determined from *k_obs_* (not carried out herein). **(B)** Simulated data for the fractions of native (dark solid lines), unfolded (light solid lines), and cleaved (dashed lines) populations over time for two proteins with equal thermodynamic stability (*K_eq_* = 10) but with either high (*k_f_* = 0.1 s^-1^, *k_u_* = 0.01 s^-1^; black) or low kinetic stability (*k_f_* = 5 s^-1^, *k_u_* = 0.5 s^-1^; red). **(C)** Image and zoomed-in schematic of SPARKflow device containing 1,104 chambers separated into 6 blocks (left). Each chamber has two compartments (DNA (green) and reaction (purple)) and 3 valves (“neck” (red), “sandwich” (orange), and “button” (blue)) (middle). Monitoring loss of fluorescence intensity over time for fluorescently-tagged and surface-immobilized proteins exposed to protease provides a high-throughput method to measure unfolding rates. **(D)** Workflow for on-chip expression, immobilization, purification, and expression quantification. **(E)** Simulated fluorescence images showing eGFP intensities after expression (left) and as a function of time after exposure to thermolysin protease (middle), yielding fluorescence intensity decay curves (right).

A variety of high-throughput methods have been developed to probe protein thermodynamic stability^22–26^, but measuring unfolding rates for many proteins in parallel has remained a challenge. Current kinetic approaches (e.g. stopped-flow kinetics^27–32^, single-molecule FRET assays^33–36^) typically explore folding pathways by monitoring changes in spectroscopic signals (*e.g.* fluorescence, circular dichroism) during chemical or thermal denaturation. However, these approaches are often limited by the time and cost associated with protein expression and purification. Labelling assays (*e.g.* hydrogen-/deuterium exchange^37–40^) can directly monitor folding and unfolding rates but require expensive instrumentation and are not easily scaled.

Quantitative native proteolysis, recently developed by the Marqusee and Park labs, presents a potentially scalable alternative^41–43^. As proteases cleave unfolded states, both thermodynamic and kinetic stability can be sensitively quantified by monitoring proteolytic cleavage over time under different conditions. For proteins that exhibit two-state folding between unfolded and native states (with folding rate *k_f_* and unfolding rate *k_u_*), *K_eq_* can be obtained from the rate of proteolysis (*k_obs_*) as the protein unfolds to a cleavable form at low protease concentrations where *k_f_* >> *k_obs_* (**Figure 1A**). More generally, for all proteins (including non-two-state proteins with folding intermediates), monitoring proteolysis under conditions where any unfolded protein is immediately cleaved (*i.e.*, cleavage is faster than refolding) reveals the rate of the first unfolding step and can be used to characterize cleavable partially unfolded forms^42,43^ (*i.e.* at high protease concentrations where *k_f_* << *k_obs_*; **Figures 1A, S1**). Gel electrophoresis provides a powerful readout of proteolysis, as resolving the size of cleavage products can also reveal structural information about unfolding intermediates^41–43^; however, these assays have been limited to tens of variants at a time.

Here, we present a novel microfluidic assay that leverages native proteolysis to enable high-throughput measurements of protein unfolding rates (SPARKfold: <u>S</u>imultaneous <u>P</u>roteolysis <u>A</u>ssay <u>R</u>evealing <u>K</u>inetics of <u>F</u>olding). SPARKfold monitors proteolytic cleavage of fluorescently-labeled, surface-immobilized proteins over time under conditions in which unfolding is rate-limiting, allowing the precise determination of 1,104 protein unfolding rate constants in less than two days. To demonstrate the power and potential of SPARKfold, we applied it to measure unfolding rates for a library of 31 DHFR orthologs and variants^43^. SPARKfold measurements agreed with those from traditional native proteolysis using both gel electrophoresis and activity-based assays; across the entire library, they revealed that single DHFR substitutions can increase unfolding rates by up to 30-fold. Comparisons with prior measurements of the impact of mutations on thermodynamic stability^43–47^ allowed us to carry out phi analysis, which suggested that the central beta sheet is disrupted in the cleavable intermediate state. SPARKfold can be applied across a wide variety of systems to quantify unfolding rates and characterize transition states, revealing biophysical principles governing protein folding pathways, informing the development of engineered proteins with enhanced kinetic stability, and uncovering how mutations drive protein misfolding and aggregation in disease.

## Results

### SPARKfold enables high-throughput expression, immobilization, purification, and native proteolysis of protein constructs

High-throughput functional characterization of large protein libraries requires the ability to recombinantly express and purify 100s-1000s of protein variants in parallel, expose expressed proteins to protease, and to follow cleavage of the expressed proteins over time. To accomplish this, SPARKfold employs microfluidic devices with 1,104 valved reaction chambers to express and immobilize fluorescently-tagged protein variants, introduce protease, and monitor loss of fluorescence due to proteolysis over time (**Figure 1C**). Each device contains 24 parallel channels that each contain 46 individual ∼1-nL reaction chambers (1104 chambers total) (**Figures 1C, S2**); chambers are grouped into 6 individually-addressable “blocks” of 4 channels each, making it possible to assay over up to 6 different conditions (*e.g.*, varying [urea], pH, salt, ligands, etc.) and perform experiments for all constructs and conditions across the device simultaneously (**Figure 1C**). Each chamber contains DNA and reaction compartments with three valves that control fluid flow: *(1)* neck valves that separate the DNA and reaction compartments, *(2)* sandwich valves that isolate reaction chambers from one another, and *(3)* button valves that enable selective surface patterning and protein purification and provide precise temporal control over when the immobilized proteins are exposed to proteases^48,49^.

To program each chamber with a unique protein variant, we align microfluidic devices to arrays of spotted plasmids that each encode a construct composed of a peptide or protein variant sandwiched between an N-terminal SNAP tag and a C-terminal monomeric eGFP tag^50^ connected via flexible glycine/serine linkers (**Figures 1D, S3**). After alignment, we actuate valves to pattern a circular surface patch within each chamber with BSA conjugated to benzyl-guanine (BG)^51^ and then introduce reagents for parallelized cell-free protein synthesis within each chamber (see Methods). After expression, the protein constructs are irreversibly recruited to patterned surfaces via a covalent bond formed between the cysteine in the active site of the N-terminal SNAP tag and BSA as the BG group is cleaved. Subsequently closing valves protects surface-immobilized constructs from flow, making it possible to wash and purify all expressed variants in parallel, and to replace the expression buffers and reagents with the conditions desired for the unfolding experiments (**Figure 1D**). After quantifying expressed variants, we introduce a protease to all chambers simultaneously and monitor the loss of eGFP fluorescence over time (**Figure 1E**). Here, we cleaved using thermolysin, a relatively nonspecific protease that preferentially cleaves hydrophobic residues expected to be exposed during unfolding. Faster unfolding results in faster loss of the native protein population and eGFP fluorescence (**Figure 1E**). Tens of devices can be fabricated in hours, and a single experiment with one device can produce up to 1104 protein unfolding curves over an additional day.

### DHFR variant library provides a test application of SPARKfold

Dihydrofolate reductase (DHFR) catalyzes the conversion of dihydrofolate to tetrahydrofolate (a key precursor in several metabolic pathways) and is a target for various small molecule therapeutics used to treat cancer and bacterial infections. *E. coli* DHFR has long served as a model system for investigating protein folding pathways^43,46,52^ and are ideal for our studies as follows: *(1)* DHFR is a small (159 amino acid), monomeric protein with no metals or disulfide bonds (**Figures 2A,B**), simplifying variant library synthesis and expression; *(2)* there are many existing datasets describing DHFR stability and unfolding rate constants to compare to our observed data^46,53^; *(3)* multiple high-resolution crystal structures (with and without bound ligands)^54,55^ facilitate interpretation of effects on residue interactions; and *(4)* multiple groups have determined that the energy landscape contains distinct kinetic intermediates that are selectively cleaved by proteases^56–63^.

**Figure 2.**
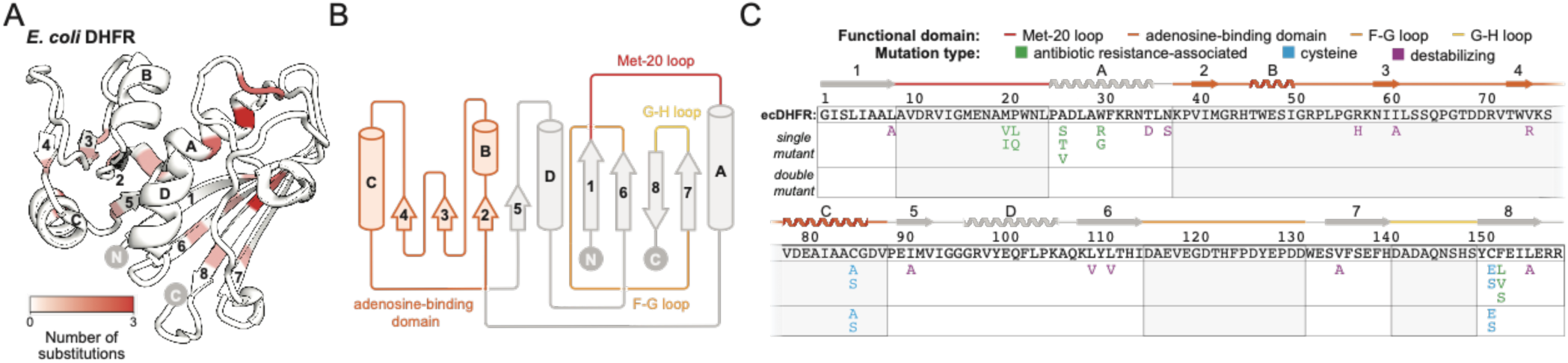
Overview of ecDHFR mutant library. **(A)** Cartoon schematic of ecDHFR structure (PDB: 6CKX) showing the sites and numbers of mutations in this study (see panel C for mutations); alpha helices (letters) and beta sheets (numbers) are labeled in ascending order from N- to C-terminal. **(B)** Cartoon schematic of topology diagram for ecDHFR structure showing locations of alpha helices (cylinders), beta sheets (arrows), Met-20 loop (red), adenosine-binding domain (dark orange), F-G loop (light orange), and G-H loop (yellow). **(C)** Schematic showing each mutation in this study along a linear topology map indicating antibiotic resistance-associated (green), cysteine (blue), or thermodynamically destabilizing (purple) substitutions; mutations and their associated references are also listed in **Supplementary Table 1**.

To develop and validate SPARKfold, we generated a library of 31 DHFR orthologs and variants: wildtype (WT) human DHFR (hDHFR), WT *E. coli* DHFR (ecDHFR), and 29 ecDHFR variants, including 12 associated with antibiotic resistance ^64–67^, 11 previously shown to be thermodynamically destabilizing^43,47,68^, and 6 with cysteine substitutions commonly used in biochemical studies and thought to have no impact on folding or activity^45^ (**Figure 2A-C; Table S1**). The ecDHFR mutations are found in alpha helices, beta sheets, and unstructured loops throughout the enzyme, including in the adenosine-binding domain and Met-20 loop that regulate catalysis^69^ (**Figures 2B,C**).

### Off-chip proteolysis for 7 constructs provides benchmark measurements and establishes unfolding is rate-limiting

Accurately quantifying protein unfolding rate constants via SPARKfold requires: (1) that protein unfolding is the rate-limiting step for cleavage, and (2) that rate constants derived from measurements of protein cleavage on-chip match those from traditional gel-based measurements^43,70^. To quantify thermolysin cleavage rates on- and off-chip, we designed positive and negative control constructs with N-terminal SNAP and C-terminal eGFP tags linked to linear peptides with flanking residues predicted to be either strongly preferred (Ala-Gly-Leu-Ala; AGLA) or disfavored (Asp-Gly-Leu-Pro; DGLP) thermolysin cleavage sites, based on prior work^71–73^ (**Figures 3A, B**). As these linear peptides do not fold, measured loss of fluorescence for these constructs should directly report on thermolysin cleavage only and quantify SPARKfold’s maximum potential dynamic range (**Figures 3A, B**).

**Figure 3.**
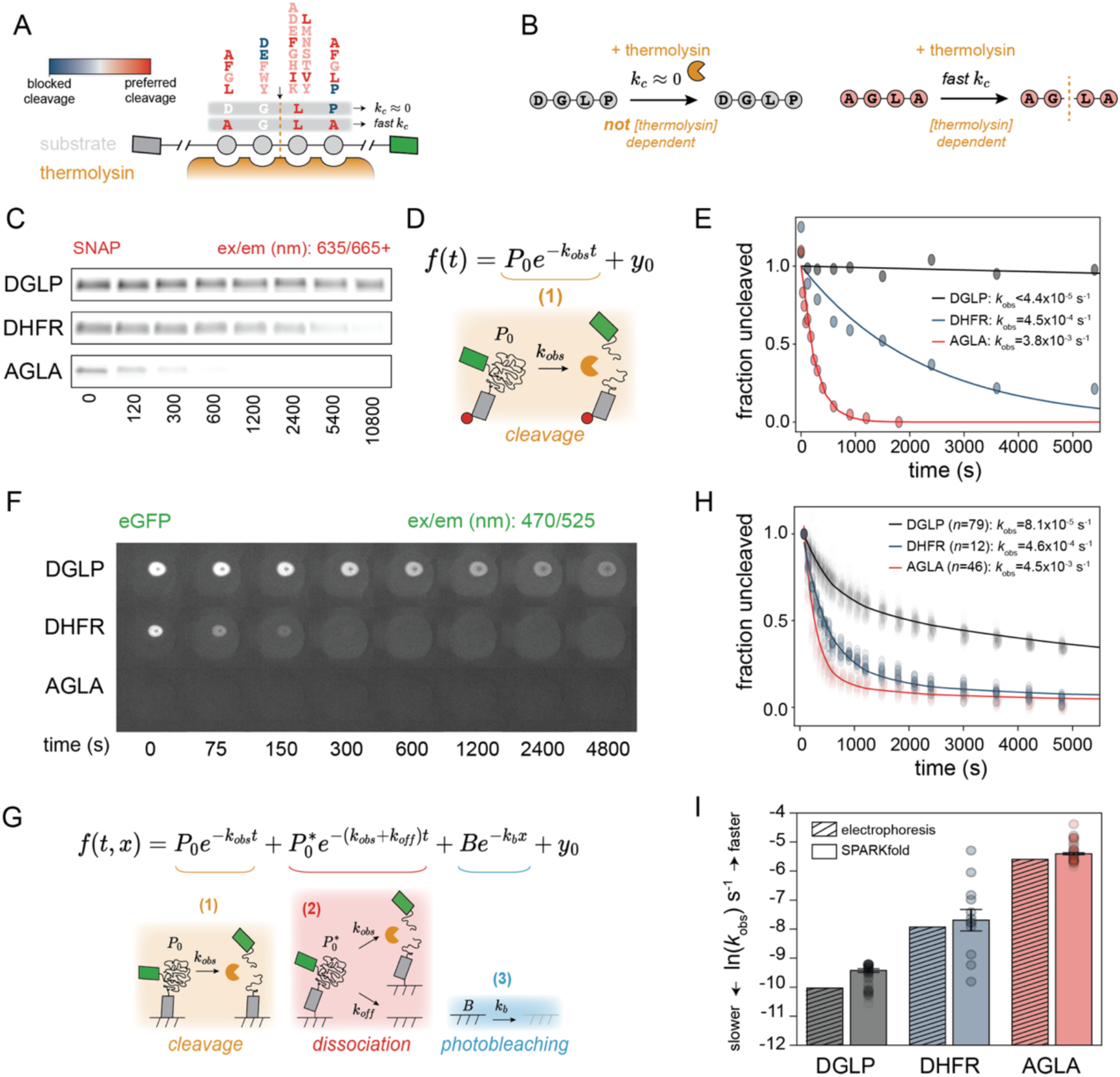
SPARKfold detects sequence-specific proteolytic cleavage and unfolding rates. **(A)** Schematic showing thermolysin (orange) specificity for residues flanking the substrate cleavage site (gray circles)^71–73^. N-terminal SNAP (gray rectangle) and C-terminal eGFP (green rectangle) tags indicate cleavage site orientation; substitutions are indicated as disfavored (blue) or preferred (red). **(B)** Negative (Asp-Gly-Leu-Pro (DGLP), left) and positive (Ala-Gly-Leu-Ala (AGLA), right) linear peptide controls for thermolysin cleavage. **(C)** Fluorescence images of SDS-PAGE gels showing amount of full-length protein constructs remaining after native proteolysis; N-terminal SNAP tag is labeled with SNAP Surface-649. **(D)** Single-exponential cleavage model for fraction of uncleaved protein (*f*) as a function of time (*t*) observed via electrophoresis; *P_0_* represents initial protein population, *k_obs_* is observed rate of proteolysis, and *y_0_* is a constant offset. **(E)** Fraction of uncleaved protein remaining as a function of time quantified from electrophoresis DGLP, WT ecDHFR, and AGLA. Markers represent normalized intensities at a given time point, lines indicate fits to single exponential decay model; measured cleavage rate for DGLP represents a lower limit. Data are normalized across multiple chambers here to show trends; intensities and fits for each chamber are available as supplemental data. **(F)** Example SPARKfold fluorescence images showing eGFP intensities as a function of time after exposure to thermolysin protease. **(G)** Composite model describing observed fraction of uncleaved protein (*f*) remaining as a function of time (*t*) and image acquisition number (*x*) that includes loss of fluorescence due to cleavage (orange), dissociation of a weakly-bound protein population (red), and photobleaching (blue). *P_0_* represents initial amount of strongly-bound protein, *k_obs_* represents observed rate of proteolysis, *P_0_\** represents initial amount of weakly-bound protein, *k_off_* represents dissociation rate, *B* is initial device and slide fluorescence, *k_p_* is photobleaching rate, and *y_0_* is a constant offset. **(H)** Fraction of uncleaved protein remaining as a function of time after SPARKfold native proteolysis for DGLP, WT ecDHFR, and AGLA. Markers represent median normalized ‘button’ intensities and lines indicate fits to plotted data using the cleavage, dissociation, and photobleaching model. For each construct, the median *k_obs_* value is reported with sample size (*n*). **(I)** Comparison between natural log-transformed *k_obs_* (ln(*k_obs_*)) values for DGLP (gray), WT ecDHFR (blue), and AGLA (red). Bars for traditional (striped) and SPARKfold (solid) measurements indicate median values with standard error (see also **Supplementary Table 2**), and scatter points display individual replicate values.

To provide benchmark off-chip measurements of unfolding rates, we used native proteolysis in solution coupled with gel electrophoresis to visualize cleavage time courses for the AGLA and DGLP positive and negative control constructs and 5 DHFR orthologs and variants with a wide range of expected unfolding rates (hDHFR, ecDHFR, and the previously characterized ecDHFR variants L8A, W30R, and V75R) ^43–47,67,68,70,74^ (**Figures 3C, S4-6, Table S2**). As expected, fluorescent band intensities for SNAP domains labeled with SNAP-Surface 649 dyes corresponding to the full-length construct decreased with increasing proteolysis time (**Figures 3C, S5**). Observed losses of intensity over time were well-fit by a single exponential decay, consistent with the expected simple Lumry-Eyring model reaction scheme^7^ (**Figures 1D, 3D-E, S6**). DGLP cleavage was slower than could be reliably detected (*k_c_* < 4.4 x 10^-5^ s^-1^), AGLA was cleaved at a much faster rate constant (*k_c_* = 3.8 x 10^-3^ s^-1^), and ecDHFR cleavage was within this range (*k_u_* = 4.5 x 10^-4^s^-1^) (**Figures 3C, 3D, S6**). To maximize assay dynamic range, these constructs (and all constructs used throughout the paper) included two mutations that removed thermolysin-preferred cleavage sites within unstructured terminal regions of the SNAP and eGFP tags to reduce rates of background cleavage (eGFP V2G/SNAP L181E) (**Figure S7**). Measurements of cleavage time courses as a function of thermolysin concentration confirmed that protein unfolding was rate-limiting: while cleavage rates for the positive control linear peptide containing a thermolysin-preferred cleavage site (AGLA) increased linearly with thermolysin concentration (*k*_c_ = 0.043(g/L)^-1^ × s ^-1^ for thermolysin concentrations ranging from 0.012 to 0.5 g/L) (**Figure S8**), cleavage rates for a ecDHFR variant with previously-measured fast unfolding rates (V75R) did not vary with increasing thermolysin (thus, unfolding is expected to be rate limiting across the library). Additional control experiments verified that thermolysin activity was not inhibited by components of the cell-free protein synthesis mixture or the presence of expressed ecDHFR constructs (**Figure S9**). Plate-based activity measurements before and after native proteolysis (**Figure S10**) yielded a loss in activity consistent with gel electrophoresis estimates of the amount of protein cleaved (*R^2^*=0.84, **Figure S11**; **Table S2**), confirming that proteolysis was degrading properly folded protein (presumably by cleaving molecules that transitioned to the unfolded state over the course of the assay).

### SPARKfold detects sequence-dependent proteolytic cleavage and returns accurate unfolding rate constants

To assess SPARKfold feasibility, we then tested whether SPARKfold could successfully express and detect sequence-specific proteolytic cleavage of surface-immobilized linear positive (AGLA) and negative (DGLP) control constructs and ecDHFR via a loss of fluorescence. As a control, we first confirmed that thermolysin activity was not inhibited by conditions required for on-chip assays, including: (1) high concentrations of BSA (above those used to prevent non-specific sticking within devices), and (2) overnight incubations in Tygon tubing at room temperature (**Figure S12**). Chambers containing plasmids encoding all 3 constructs showed substantial eGFP intensities while empty chambers did not, confirming successful expression and surface-immobilization of SNAP and eGFP-tagged proteins without cross-contamination (**Figure 3F**). As expected, fluorescence images revealed that eGFP fluorescence was lost fastest for the positive control (AGLA) construct, consistent with sequence-specific proteolysis of surface-immobilized protein (**Figure 3F**). However, while solution assays of DGLP revealed minimal cleavage, chambers containing this construct showed a considerable decrease in eGFP fluorescence over time (**Figure 3F**). This behavior was well fit by a functional form derived for a reaction scheme that includes photobleaching (proportional to the number of times the sample is exposed to light, see below) and two populations of surface-immobilized molecules: (1) a strongly-bound population that is cleaved by thermolysin and (2) a weakly-bound population that can either be cleaved or slowly dissociate from the surface (**Figures 3G,H**; see Supplementary Methods for derivation).

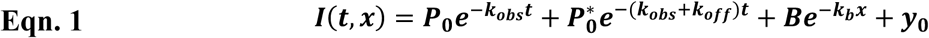

Here, *P*_0_ is the starting population of strongly-bound molecules, *k*_obs_ is the observed cleavage rate constant, *P*\**_0_* is the starting population of weakly-bound molecules, *k*_off_ represents their dissociation rate constant, *B* is the initial fluorescence of molecules that photobleach over the course of the experiment, *k*_b_ is the photobleaching rate constant, *t* is time, x is the number of images acquired at a given timepoint, and *y*_0_ represents an initial constant offset. Each SPARKfold experiment includes empty and DGLP-containing chambers throughout, making it possible to quantify photobleaching and slow dissociation rate constants independently. We therefore determined observed cleavage rates for DHFR variants via a 3-step process. First, we determined a median amplitude and rate constant for photobleaching by fitting observed intensity *vs.* image number from all blank chambers to a single-exponential decay (**Eqn. 2, Figure S13**):

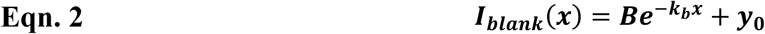

Next, we determined a median rate constant for slow dissociation by fitting observed intensity *vs.* time from negative cleavage control (DGLP)-containing chambers with the photobleaching rate held constant and assuming *k*_off_ >> *k*_obs_ (as for a non-cleavable substrate; **Eqn. 3, Figure S14**):

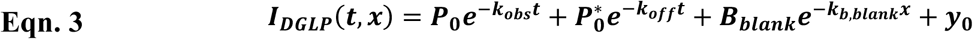

Finally, we fit observed rate constants for cleavage (and thus inferred unfolding rate constants assuming unfolding is rate-limiting) for AGLA- and ecDHFR-containing chambers using **Eqn. 1** and the median photobleaching and slow dissociation rate constants obtained above (**Figures 3H, S15-16**):

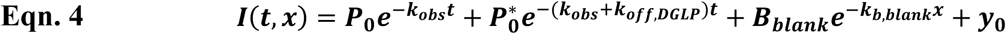

With this procedure, median SPARKfold-returned cleavage rate constants across all chambers for DGLP, WT ecDHFR, and ALGA agreed with those determined by traditional gel-based native proteolysis within 2-fold over a 150-fold range of *k*_obs_ values (**Figure 3I**).

### SPARKfold enables parallel unfolding rate measurements for the DHFR variant library

Next, we applied SPARKfold to quantify unfolding rates for all 31 orthologs and variants within the DHFR library, each at high redundancy to ensure excellent statistics. Chambers containing ecDHFR variants demonstrated higher expression and surface immobilization compared to empty chambers, although these levels were significantly lower than for chambers with control peptide constructs (**Figure 4A**). Based on these distributions, we set a minimum intensity threshold to exclude chambers with poor expression from downstream analysis (**Figure 4B**), yielding 90% of DHFR orthologs and variants (28/31) with 5 or more chambers passing this threshold across experiments (**Figure 4B**). To estimate unfolding rate constants for each variant, we again: (1) quantified rate constants for photobleaching from blank chambers (**Figure S13**), (2) used these photobleaching rate constants and data from DGLP-containing chambers to quantify surface dissociation rate constants (**Figure S14**), and then (3) fit observed fluorescence over time to a kinetic model that includes these processes and unfolding (**Figures S16-17**). Raw curves for every chamber within each of 2 replicate experiments and compiled summary data across each experiment (**Figure S18**) are available as Supplementary Files and in an associated OSF repository^75^. Median cleavage rates (for control peptides) and unfolding rates (for DHFR variants) replicated reasonably well across experiments (*R^2^* =0.84, **Figure 4C**; **Table S3**) and agreed with those determined with gel-based native proteolysis (**Figure 4D**, *R^2^* =0.95). In general, constructs with relatively low initial eGFP intensities yielded rate constant distributions with the highest variance (**Figure 4E**), suggesting that future improvements of on-chip immobilization could improve assay precision (see Discussion).

**Figure 4.**
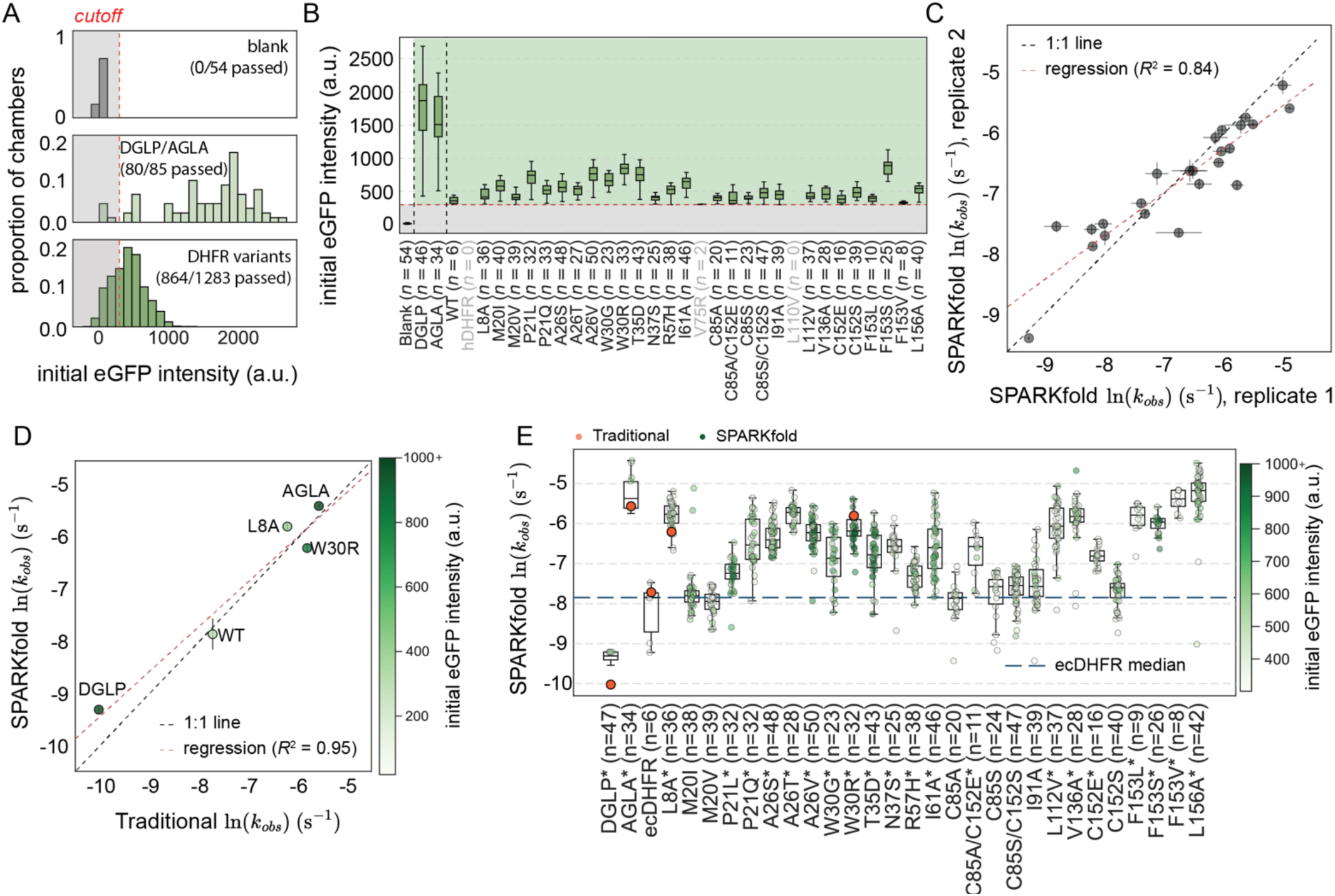
SPARKfold expression and unfolding rate measurement for library variants. **(A)** Histograms of per-chamber initial background-subtracted eGFP intensities for blank (top), control peptide (DGLP/AGLA) (middle), and DHFR mutant (bottom) chambers; annotation indicates sample size (*n*) for each. Red dashed line represents a threshold of 156 a.u. to filter out chambers with low expression. Each histogram is normalized to show the proportion of data within each bin. **(B)** Boxplot of initial background-subtracted eGFP chamber intensities by construct. Boxes span the interquartile range (IQR), lines inside boxes represent the median, and whiskers extend to 1.5xIQR beyond the box. Red dashed line marks the expression threshold (156 a.u.); shading distinguishes blank chambers (left), control peptide chambers (DGLP/AGLA) (middle), and protein chambers (DHFR mutants) (right). Construct labels include sample sizes (*n*). **(C)** Scatter plot comparing natural log-transformed *k_obs_* (ln(*k_obs_*)) values for two SPARKfold experimental replicates. Each marker represents a construct, error bars indicate the standard error of the median ln(*k_obs_*) for each replicate, red dashed line indicates linear fit to plotted data (*R^2^* = 0.87), and black dotted line indicates the 1:1 relationship. **(D)** Scatter plot comparing ln(*k_obs_*) values for SPARKfold and traditional gel-based native proteolysis experiments across replicates. Each marker represents a construct, error bars indicate per-replicate standard error of the median ln(*k_obs_*), red dashed line indicates linear fit to plotted data (*R^2^* = 0.87), and black dotted line indicates the 1:1 relationship. Gradient color bar reflects the median initial background-subtracted eGFP intensity with higher values capped at 1000, illustrating that experimental variance is highest for variants with low expression. **(E)** Boxplot with overlaid strip plot comparing ln(*k_obs_*) values for SPARKfold and traditional gel-based native proteolysis experiments by construct. Boxes span the interquartile range (IQR), lines inside boxes represent the median, whiskers extend to 1.5xIQR beyond the box, and points outside this range are shown as outliers. Stripplot markers (green) display individual replicate values. Gradient color bar reflects the median initial background-subtracted eGFP intensity, with higher values capped at 1000. Markers (orange) show ln(*k_obs_*) values from traditional gel-based experiments, and blue dashed line indicates median ln(*k_obs_*) for WT ecDHFR. Construct labels include sample sizes (*n*) and Bonferroni-corrected significance markers (* for *p* < 0.05) compared to WT ecDHFR; values are also given in **Supplementary Table 3**.

Measured unfolding rate constants across ecDHFR variants varied by ∼20-fold. Most variants (17/28) unfolded significantly faster than WT and no variants unfolded more slowly (as determined by bootstrapped significance estimation with Bonferroni correction for multiple comparisons; **Figures 4E, S19; Table S3**). Proteolysis rates for the WT and 7 variants of ecDHFR were previously measured via traditional native proteolysis with a gel electrophoresis readout^43^; however, these initial measurements used a lower concentration of thermolysin (to probe the equilibrium between the native and cleavable intermediate states rather than unfolding rate constants) and did not consider possible impacts of product inhibition. SPARKfold-returned measurements show similar sequence-dependent trends and agree with published rates when corrected for expected product inhibition^70^ (**Figure S20**).

### Substituting charged residues within the hydrophobic core drives large increases in unfolding rate

The suite of unfolding rates returned by SPARKfold provides a new opportunity to search for sequence and structural features that shape the DHFR unfolding pathway. Projecting SPARKfold-measured unfolding rates onto the ecDHFR structure revealed no simple relationship between the spatial distribution of variants and their impact on kinetic stability (**Figure 5A**). Consistent with expected impacts of disrupting residue packing interactions and secondary structure formation, mutations to residues within beta sheets, alpha helices, and unstructured loops increased median *k_u_* by ∼6-fold, 3-fold, and 2-fold, respectively (**Figure 5B**). Nevertheless, the magnitude of observed effects within secondary structure elements varied substantially depending on the tertiary contacts made within the native structure and the properties of the substituted residue (**Figures 5A and 5B**). Disrupting beta sheet residues that directly contact other residues in the native structure generally increased *k_u_* (*e.g.* L8, L112, V136, and L156, which pack against one another to form an anti-parallel sheet and hydrophobic core (**Figures 5A(iii) and 5B**); F153, which packs and forms pi-pi stacking interactions with A26 and W30 on an alpha helix (**Figures 5A(iv) and 5B**)). At a given residue, the magnitude of the effect varied with the magnitude of the biochemical perturbation (*e.g.* substituting a cysteine with a charged residue (C152E) was >2-fold more deleterious than a more conservative mutation (C152S)) (**Figures 5A(iv) and 5B**). To test if measured unfolding rates for mutations in alpha helices could be explained by changes in helical propensity, we compared measured changes in *k_u_* with previously determined changes in Gibbs free energy for helix formation^76^ (see Methods). For mutations within alpha helices, predicted changes in helical propensity accounted for ∼50% of observed increases in unfolding rates (**Figure 5C**). The W30R mutation increases *k_u_* more than predicted by the change in helical propensity, likely due to disrupted interactions with the beta sheet residue F153 (**Figures 5A(iv) and 5C**). Most residues within unstructured regions are found on the Met-20 loop. While M20 mutations do not alter *k_u_*, consistent with expectations for a flexible region, P21 substitutions on the same loop raise *k_u_* by ∼2-4-fold, potentially by enabling greater conformational flexibility (**Figure 5B**). Any insertion of a charged residue within the buried hydrophobic core increased unfolding rates by >2-fold (**Figures 5D-E, S21**).

**Figure 5.**
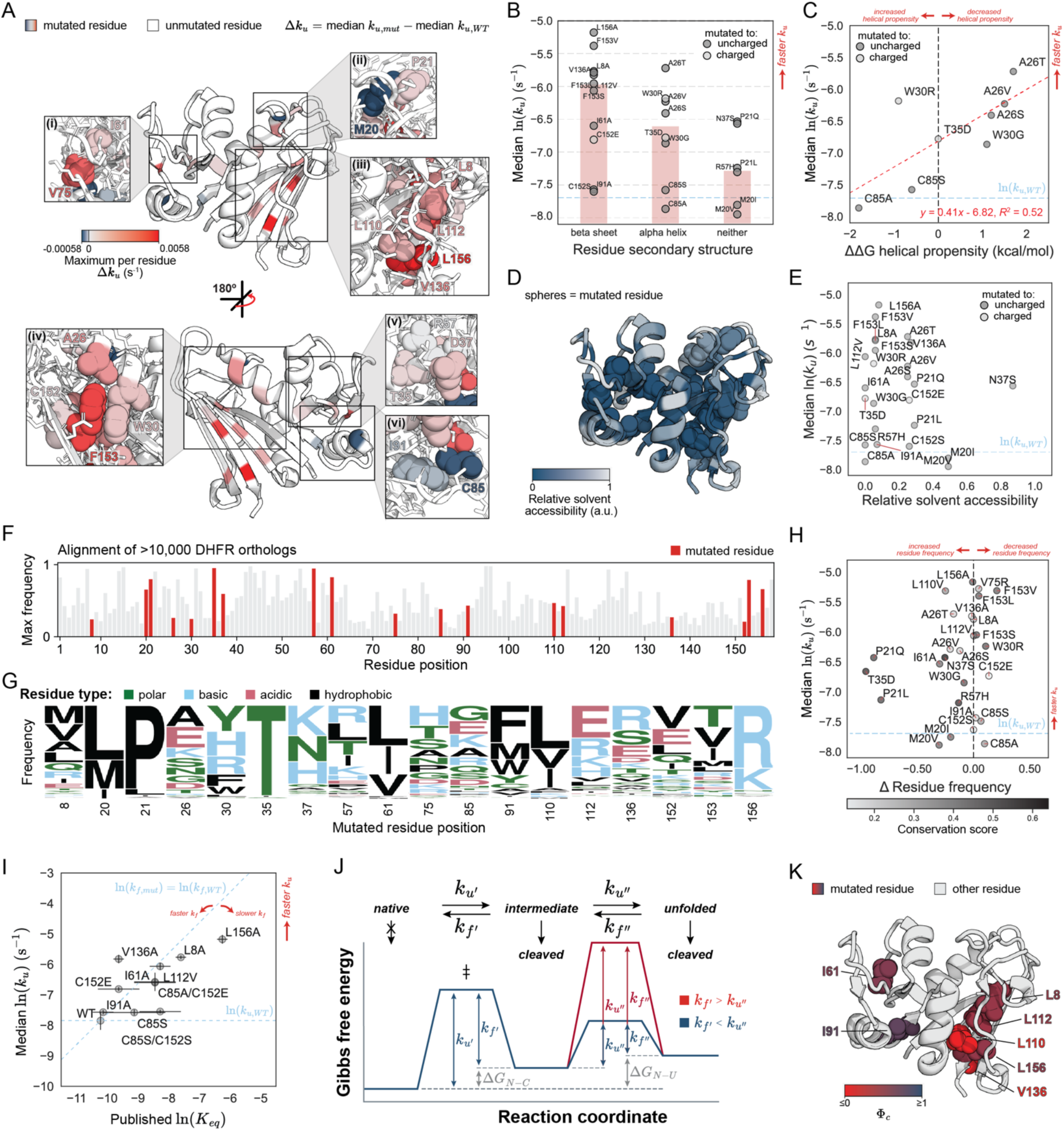
Analysis of ecDHFR unfolding rates based on structural and evolutionary features. **(A)** Entire and zoomed-in portions of the structure (PDB: 6CXK) showing the difference in measured unfolding rate between the mutant and WT ecDHFR (Δ*k_u_*) for the substitution with the largest effect at that position. Residue color ranges from no significant effect (Δ*k_u_* < 0; blue) to increased (Δ*k_u_* > 0; red) unfolding rates relative to WT; at mutated positions, the native residue is displayed as spheres. **(B)** Bar plot and overlaid stripplot comparing natural log-transformed *k_u_* (ln(*k_u_*)) values for each construct based on secondary structure at substitution position. Stripplot markers are colored based on mutation to an uncharged (filled circle) or charged (empty circle) residue. Bar heights indicate median ln(*k_obs_*) for substitutions made in beta sheets (left), alpha helices (middle), and neither (right). Blue dashed line indicates median ln(*k_u_*) for WT ecDHFR. **(C)** Scatter plot comparing median ln(*k_u_*) values for ecDHFR variants in alpha helices with previously determined changes in Gibbs free energy for helix formation, ΔΔ*G* helical propensity (kcal/mol)^76^. Markers are colored based on mutation to an uncharged (filled circle) or charged (empty circle) residue. Red dashed line indicates linear fit to plotted data (*R^2^* = 0.54), blue dashed line indicates median ln(*k_u_*) for WT ecDHFR, and black dashed line indicates no change in helical propensity (ΔΔ*G* = 0 kcal/mol). **(D)** Relative solvent accessibility of residues on the ecDHFR structure with residues mutated in this study shown as spheres. All ecDHFR residues are colored on a scale from low (dark blue) to high (light gray) relative solvent accessibility. **(E)** Scatter plot comparing median ln(*k_u_*) values for ecDHFR variants with relative solvent accessibility. Markers are colored based on mutation to an uncharged (filled circle) or charged (empty circle) residue. Blue dashed line indicates median ln(*k_u_*) for WT ecDHFR. **(F)** Frequency of the most common residue at each position calculated from a multiple sequence alignment of >10,000 DHFR orthologs; red bars indicate mutated residue positions. **(G)** Relative residue frequencies at each mutated residue positions colored based on residue type: polar (green), basic (blue), acidic (red), and hydrophobic (black). **(H)** Scatter plot comparing median ln(*k_u_*) values for ecDHFR variants with change in frequency of the substituted relative to the native residue. Markers are colored based on conservation at the mutated residue position, ranging from less conserved (white) to more conserved (black). Blue dashed line indicates median ln(*k_u_*) for WT ecDHFR and black dashed line indicates no change in residue frequency. **(I)** Scatter plot comparing median ln(*k_u_*) values for ecDHFR variants with ln(*K_eq_*), calculated from published free energy of unfolding (Δ*G_u_*) values^43–47,68^. Vertical error bars indicate the standard error of the median ln(*k_obs_*) for each replicate; horizontal error bars indicate the lower and upper limit of ln(*K_eq_*) calculated from the range of previously determined Δ*G_u_* values; blue dashed line indicates median ln(*k_u_*) for WT ecDHFR. Dashed diagonal line shows the expected correlation between *k_u_* and *K_eq_* if stability is only dependent on the unfolding rate; mutants to the left of the dashed line have a faster *k_f_*, while mutants to the right have a slower *k_f_* relative to WT ecDHFR. **(J)** Gibbs free energy *vs.* reaction coordinate diagram for native, intermediate, and unfolded protein populations, where intermediate and unfolded proteins can be cleaved by protease. Height of the energy barrier between the native and intermediate states depends on the unfolding rate (*k_u’_*) and folding rate (*k_f’_*); height of the energy barrier between the intermediate and unfolded states depends on the unfolding rate (*k_u’’_*) and folding rate (*k_f’’_*). Change in Gibbs free energy (gray) is annotated between native and cleavable states (Δ*G_N-C_*) and between native and unfolded states (Δ*G_N-U_*). Example diagrams shown under conditions where *k_f’_* > *k_u’’_* (red) and *k_f’_* < *k_u’’_* (blue). **(K)** Calculated phi (Φ_c_) values displayed on the ecDHFR structure (PDB: 6CXK). Mutated residues with thermodynamic stability data available to calculate Φ_c_ are displayed as spheres and colored from Φ_c_ ≤ 0 (red) to Φ_c_ ≥ 1 (blue) (see also **Supplementary Table 4**).

### Evolutionary conservation does not correlate with observed impacts on unfolding rates

Enzymes must fold to function. Prior studies have reported correlations between mutational effects on stability and residue conservation within phylogenetic alignments and suggested the existence of a stability ‘threshold’ below which fitness is compromised^77–84^. However, it remains unclear whether such ‘thresholds’ are dictated by thermodynamic stability, kinetic stability, both, or other factors. To investigate the relationship between evolutionary conservation and the impacts of mutations on folding rates, we aligned >10,000 DHFR orthologs and quantified residue frequencies at each position (**Figures 5F and 5G**). Impacts on *k*_u_ were largely uncorrelated with conservation, with many mutations to equally probable residues yielding substantial increases (**Figure 5H**). While several previous studies have demonstrated that ecDHFR mutational landscapes vary substantially upon deletion of the quality control protease Lon ^85,86^, we find no significant correlation when comparing either measured *k*_u_ or published *K_eq_* values with Lon selection coefficients (**Figure S22**). These results underscore the complexity of biological selection. While these results are consistent with prior DHFR studies that suggest fitness depends on a competition between chaperonins and proteases for access to unfolded intermediates^85^, establishing this *in vitro* would require additional measurements of rate constants in the presence and absence of different proteases and chaperones.

### Most DHFR variants reduce thermodynamic stability primarily by increasing the rate of unfolding

To gain insight into the DHFR unfolding pathway, we compared measured impacts of mutations on unfolding kinetics with previously-published equilibrium stability measurements (**Figure 5Ι**). For proteins that follow a two-state folding model, *K*_eq_ = *k*_f_/*k*_u_ such that mutations that cause *<u>the same</u>* change in *K_eq_* and *k_u_* (*i.e.* on diagonal line, **Figure 5I**) can be inferred to have no impact on *k_f_*. However, for proteins with intermediates, such as ribonuclease H (RNase H), thermodynamics and protease cleavage-derived kinetics can be decoupled^87^. The DHFR folding pathway contains cleavable intermediate and unfolded states^46^ such that measured proteolysis rates may either report on the kinetics of partial unfolding (*k_u’_*) or global unfolding (*k_u’’_*), depending on which step is rate-limiting and which states are cleavable (**Figure 5J**); thus, thermodynamic stability and measured proteolysis rate cannot be used to reliably infer the folding rate. Two mutations that pack against one another within the beta sheet (V136A and L156A) increased apparent unfolding rate constants (**Figures 5I and 5K**); while V136A slightly decreased thermodynamic stability compared to WT, L156A decreased thermodynamic and kinetic stability similarly (**Figures 5I and 5K**).

By comparing published *K_eq_* and measured *k_u_* values, we can determine phi (Φ_c_), the ratio in the log-transformed mutation-induced change in the unfolding rate constant to the change in thermodynamic stability of the native form, thereby probing structures of the transition state intermediates along the unfolding pathway. Mutating residues that are unstructured within the unfolding transition (*i.e.* cleavable, high-energy) state alters the unfolding rate and stability of the native forms equally (Φ_c_ = 1), while mutating residues that are structured in this transition state affect the stability of the native form alone but does not change the kinetic barrier (Φ_c_ = 0; **Figure 5K**, see Methods and **Table S4**). Projecting Φ_c_ values onto the structure suggests that the beta sheet is primarily unfolded in the transition state while the adenosine binding domain remains folded (**Figure 5K**), in agreement with previous conclusions based on the thermodynamic stability of the cleavable intermediate (Δ*G_N-C_*)^43^.

## Discussion

Most published studies of protein stability focus on thermodynamic stability, likely because thermodynamic stability is more easily measured and predicted by existing assays and algorithms^11^. Yet in many cases, kinetic stability can be equally or more important in determining function. Secreted proteins that must function in harsh extracellular environments are often kinetically hyperstable, limiting their risk of degradation^5^. Within cells, partial or complete unfolding can regulate signal transduction by modulating effective ligand binding affinities and/or accessibility of regions subject to post-translational modifications^88^. Finally, proteins used in industrial or biotechnological applications face harsh solvent conditions and high risk of irreversible denaturation^11^. Here, we present a novel technology (SPARKfold) that can be used to quantify kinetic stability under native (non-perturbative) conditions at scale.

Previous measurements of equilibrium constants (*K_eq_*) and unfolding rates (*k_u_*) for ∼30 single-domain two-state proteins varied by 7 orders of magnitude^89^. Here, most amino acid substitutions within ecDHFR increased unfolding rates by <2-fold, with the most deleterious substitutions increasing unfolding rates by ∼20-fold. As mutations that dramatically alter kinetic stabilities would likely be subjected to strong negative selection, the small dynamic range observed here likely stems from our decision to profile variants observed in wild populations. Consistent with this model, several mutations with only minor impacts on unfolding rates were previously shown to significantly affect global stability and ligand binding (*e.g.* trimethoprim resistant and destabilized ecDHFR variants P21L and A26T^67,90^).

DHFR tends to aggregate under the low-pH conditions required for hydrogen/deuterium exchange assays and facilitate site-specific introduction of chemical modifications using thiol-reactive probes^91^. To prevent this aggregation, several prior studies have investigated cysteine-free double mutant variants^45,92–94^ thought to retain similar activity, thermodynamic stability, and folding mechanism to the WT protein^45^. Here, we find that the single and double cysteine mutations to alanine and serine have no statistically significant impact on unfolding rates, confirming the utility of these constructs for future studies of DHFR kinetic stability. The commonly used C152E mutation^93,95,96^ shows a small yet significant increase in unfolding rate (∼3-fold, p<10^-6^; potentially due to a change in surface charge), suggesting that C152S should be used instead.

SPARKfold is compatible with any protein that: *(1)* is not susceptible to proteolysis in the native state, *(2)* exposes a protease-compatible cleavage site when unfolding, and *(3)* refolds more slowly than the rate of proteolysis at feasible protease concentrations. While we used thermolysin for all measurements here, additional measurements using proteases with differing specificities could provide additional information about local unfolding intermediates. If the globally unfolded state is the lowest-energy cleavable state, SPARKfold can provide quantitative information about global unfolding rates; otherwise, SPARKfold provides information about local unfolding. Here, the fastest and slowest unfolding rates that could be resolved were *k_upper_*=0.013s^-1^ and *k_lower_*=8.1 x 10^-5^ s^-1^. In future work, the upper limit of this dynamic range could be increased by altering the imaging setup to enable higher-throughput image acquisition. The lower limit of this dynamic range was imposed by observed slow dissociation of surface-immobilized constructs similar to DGLP cleavage rates observed off-chip, suggesting cleavage of either the SNAP or eGFP tags, the flexible linkers, or the BSA adsorbed to the device surface. As variance in measured unfolding rates was inversely correlated with expression levels below a certain threshold, future optimization of variant DNA preparation, library printing, and on-chip expression conditions could boost expression and reduce this variability.

Beyond measuring unfolding kinetics, SPARKfold lays the foundation for a wide variety of additional assays. SPARKfold enables precise measurement of small changes in *k_u_* for many mutations at each residue position without disrupting the global structure. Thus, SPARKfold measurements can reveal mechanisms by which intermediate kinetics may impact a protein’s cellular abundance or propensity to aggregate; measurements monitoring cleavage by multiple proteases with different cleavage preferences could further provide clues regarding intermediate structures. When combined with *K_eq_* measurements, SPARKfold provides unique insights into folding intermediates suggested to be conserved across orthologs for complex enzymes^87^. As ligand binding stabilizes protein folded conformations, comparing unfolding rates measured in the presence and absence of drug could provide a sensitive method for high-throughput quantification of binding of unlabeled ligand. Finally, applying SPARKfold to varying linear (unfolded) sequences could provide a high-throughput method for systematically investigating the determinants of protease specificity.

## Resource Availability

### Lead Contact

Further information and requests for resources and reagents should be directed to and will be fulfilled by the Lead Contact, Polly Fordyce.

### Material availability

Plasmids generated in this study have been deposited to AddGene.

### Data and Code Availability

Raw data from all unfolding measurements has been deposited at Open Science Framework (https://doi.org/10.17605/OSF.IO/EGCU4). Code and input files used to analyze data and generate the figures reported in this paper are available under the “AnalysisScriptsAndRequiredInputs” folder; supplementary data files summarizing results per experiment and across experiments are available in the “SupplementalFiles” folder. Automation software associated with experimental and microscopy setups is located at https://pypi.org/project/acqpack/ and https://github.com/FordyceLab/RunPack. Software for processing images is available at https://github.com/FordyceLab/ImageStitcher. Analytical software is available at https://github.com/FordyceLab/ProcessingPack. Use of other software is described in the STAR Methods section. Any additional information required to reproduce this work is available from the Lead Contact.

## Acknowledgements

This work was supported by NIH Grant number R01GM064798 awarded jointly to D.H. and P.M.F.; P.M.F. is also a Chan Zuckerberg Biohub San Francisco Investigator. B.A. was supported by an NSF Graduate Research Fellowship. The authors thank all members of the Fordyce and Herschlag labs for helpful feedback on the manuscript. In particular, we thank Professor Susan Marqusee and Shawn Costello for sharing their vast knowledge about protein folding and stability, Karl Krauth for his insights into mathematical modeling, Kara Brower for her help with microfluidic device design, and Matt DeJong for his conceptual help with protein surface immobilization.

## Author Contributions

Conceptualization, all authors; investigation, B.A. and F.S.; writing – original draft, all authors; writing – review and editing, all authors; supervision and funding acquisition, P.M.F and D.H.

## STAR METHODS

### Key resources table

| Reagent or resource | Source | Identifier |
| --- | --- | --- |
| **Bacterial and Virus Strains** |  |  |
| NEB 5-alpha Competent *E. coli* | New England Biolabs | C2987I |
| **Chemicals, Peptides and Recombinant Proteins** |  |  |
| UltraPure BSA | ThermoFisher | AM2618 |
| Phosphate Buffered Saline (10X) | Corning | 46-013-CM |
| Trehalose Dihydrate | Sigma Life Science | T9531 |
| Sodium Chloride | Sigma | 204439 |
| NHS-GLA-BG | New England Biolabs | S9151S |
| Thermolysin | Sigma | T7902 |
| Borate buffer | LabChem | LC117001 |
| EDTA | Ambion | AM9260G |
| SNAP-Surface 649 | New England Biolabs | S9159S |
| Abz-AGLA-Nba | MedChemExpress | HY-P4406 |
| Poly-L-lysine | Electron Microscopy Sciences | 19320-A |
| Dihydrofolic acid | Sigma Aldrich | D7006 |
| NADPH | Thermo Fischer | AC328742500 |
| Calcium Chloride | Sigma | C2536 |
| **Critical Commercial Assays** |  |  |
| PURExpress *In Vitro* Protein Synthesis Kit | New England Biolabs | E6800L |
| RNasin Ribonuclease Inhibitor | Promega | N2515 |
| DpnI | New England Biolabs | R0176L |
| QIA Prep spin miniprep kit | QiaGen | 27104 |
| **Deposited Data** |  |  |
| Raw unfolding measurements, analytical code | This paper | https://doi.org/10.17605/OSF.IO/NAWME |
| **Software and Algorithms** |  |  |
| AcqPack | PyPI | https://pypi.org/project/acqpack/ |
| ImageStitcher | GitHub | https://github.com/FordyceLab/ |
| ProcessingPack | GitHub | https://github.com/FordyceLab/ProcessingPack |
| RunPack | GitHub | https://github.com/FordyceLab/RunPack |

### Methods details

#### Cloning DHFR into PURExpress vector for cell free expression

We cloned the gene coding for *E. coli* DHFR (ecDHFR), a thermolysin-preferred substrate (Ala-Gly-Leu-Ala; AGLA), and a thermolysin-resistant substrate (Asp-Gly-Leu-Pro; DGLP) into the PURExpress plasmid, which contains a C-terminal eGFP tag and an N-terminal SNAP tag, both attached with a flexible glycine/serine linker (GGGSGGG). We confirmed the sequence of the construct with Sanger sequencing, and all subsequent mutagenesis used the ecDHFR construct sequence as a plasmid template.

#### Optimization of the tags

To remove alternative expression start sites and stabilize the construct against proteolysis outside of the protein of interest, we used PeptideCutter to identify thermolysin cleavage sites in the unstructured regions between the protein and both the eGFP and SNAP tags. We performed site-directed mutagenesis (as described below) to generate the following mutations for all constructs: SNAP L181E and eGFP M1G/V2G. For the DHFR construct, we introduced an additional mutation to remove the start codon (DHFR M1G). After confirming these optimized construct sequences with Sanger sequencing^75^, we used the two control constructs for thermolysin (SNAP-AGLA-eGFP and SNAP-DGLP-eGFP) to test the proteolysis resistance of the construct with and without stabilizing mutations, following rates of native proteolysis (described above).

#### Site-directed mutagenesis

We performed site-directed mutagenesis using the standard protocol for Pfu Turbo. We used the PCR product to transform chemically competent DH5a cells, which we made in-house following standard protocols, and plated them on LB agar carbenicillin plates. We picked single colonies and grew them in 10 mL LB with 50 µg/mL carbenicillin overnight on a rotor at 37°C. After pelleting the cells, we purified the plasmids using the GeneJet Plasmid Miniprep Kit (Thermo Scientific) and verified the sequence of the genes with Sanger sequencing.

#### Cell free expression

We used the following protocol for the expression of double-tagged DHFR with the PURExpress *in vitro* synthesis kit (NEB): (1) we incubated 30 µL of Solution A with 22.5 µL of Solution B on ice for 45 minutes, (2) we added 1.5 µL of RNAsin, 0.75 µL of 100 mM DTT, 19 µL of UltraPure water, and 1 µL of ∼200 ng/µL plasmid, and then (3) we ran the *in vitro* transcription-translation reaction in a PCR machine for 2 hours at 30°C followed by 90 minutes at 23°C. After expression, we added 75 µL of 100 mM Tris•HCl (pH 8.0), 150 mM NaCl for unlabeled SNAP, and to label SNAP we added 10 nM benzyl guanine (BG)-conjugated Dynomics dye DY-649P1 (SNAP-Surface 649, NEB) and incubated at room temperature for 1 hour. We measured the amount of expressed protein based on eGFP fluorescence using a Denovix and calculated the concentration using an eGFP standard curve. For off-chip experiments, we buffer-exchanged the PURExpress in 0.5 mL Amicon filters (Sigma) 3 times with 100 mM Tris (pH = 8), 150 mM NaCl, and 1 mM DTT.

#### Preparation of thermolysin

We prepared 10 g/L thermolysin from *Geobacillus stearothermophilus* (Sigma) as described by Marqusee and Park^97^. We dissolved 100 mg of thermolysin in 10 mL of 2.5 M NaCl and 100 mM CaCl_2_, separated the solution into 50 200-µL aliquots, and stored at −20°C.

#### Test of loss of activity of thermolysin over time

To assess if the observed rate of proteolysis was influenced by a change in the concentration of active thermolysin over time, we used a commercial protease substrate abz-AGLA-nBa (MedChemExpress) to measure the activity of thermolysin after incubation on the SPARKfold device, with BSA, or with PURExpress components. We dissolved the substrate in 30% acetic acid and stored it at −20°C. We incubated thermolysin (prepared as described above) in Tygon tubing (Saint-Gobain, AAD04103), BSA over a 10^5^-fold range (0.15 nM-15 µM), or with WT or V75R ecDHFR in PURExpress overnight. We then observed turnover kinetics in 384 plates at 314/417 nm on a Tecan M200, using 100 mM Tri•HCls (pH 8.0), 150 mM NaCl and 0.3–0.8 µM substrate.

#### Native proteolysis, gel-based assay

To determine if the observed rate of proteolysis was limited by the *K_eq_* between folded and unfolded protein or by the unfolding rate, we quantified the rate of DHFR proteolysis across a 20-fold range of thermolysin concentrations (0.06-0.5 g/L). We buffer-exchanged DHFR expressed with PURExpress into 100 mM Tris•HCl (pH 8.0) and 150 mM NaCl in 3K Amicon filters, diluting the buffer 100-fold. To the buffer-exchanged DHFR, we added different concentrations of thermolysin and collected timepoints by quenching 4 µL aliquots with 4 µL of 50 mM EDTA. We stored the aliquots on ice under tin foil until all time points had been collected. To each sample, we added 3 µL of Laemmli loading dye, ran the samples on denaturing gels (150V, 45 minutes), and imaged gels on a Typhoon FLA 9500 (for GFP: excitation blue laser (473 nm) and emission (>510 nm); for SNAP649: excitation red laser (635 nm) and emission (>665 nm)) or BioRad Versadoc. We used ImageJ to quantify the intensity of the gel bands corresponding to the full-length construct, plotted the concentration of full-length construct as a function of proteolysis time, and fit the data to a single exponential function. To ensure that observations that unfolding is rate-limiting for DHFR were not due to solubility issues or similar effects that would result in non-first order thermolysin cleavage kinetics, we also performed the above experiment with thermolysin substrate AGLA, varying the thermolysin concentration over a 100-fold range (0.012-1.0 g/L).

#### Native proteolysis, activity-based assay

To provide an orthogonal readout of thermolysin cleavage rates (and verify that thermolysin was cleaving active protein that had unfolded), we quantified enzymatic activity after native proteolysis at multiple timepoints. First, we exposed DHFR constructs to thermolysin and collected timepoints as described for the denaturing gel assay above. We then added 4 µL of each DHFR sample to 200 µL of 100 mM Tris•HCl (pH 8.0), 150 mM NaCl, 1 mM DTT, 50 µM dihydrofolic acid (Sigma Aldrich D7006-10MG) and 60 µM NADPH (Thermo Fischer AC328742500) in a 96-well plate. Using a Tecan M200, we measured the DHFR activity in each sample by monitoring the change in substrate concentration at 340 nm, corresponding to conversion of substrate NADPH to product NADP+ by the active enzyme. For each sample, we fit observed fluorescence *vs.* proteolysis time to a single exponential decay model.

#### Mold and device fabrication

Flow and control molding masters were fabricated as previously described^98,99^. Using these molds, we cast two-layer microfluidic devices using polydimethylsiloxane (PDMS) polymer (RS Hughes, RTV615). To generate control layers (**Figure S1**, orange), we combined 60 grams of PDMS components (1:5 ratio of cross-linker to base), mixed with a THINKY centrifugal mixer for 3 minutes at 2000 rpm, poured the mixture onto the molds, degassed in a vacuum chamber for 45 minutes, and then baked in a convection oven for 50 minutes at 80°C. After cutting and removing device control layers from the wafer, we added control inlets using a hole punch (SYNEO, CR0350255N20R4) mounted on a drill press (Technical Innovations). To generate flow layers (**Figure S1**, blue), we combined PDMS components (1:20 ratio of cross-linker to base), mixed with a THINKY centrifugal mixer for 3 minutes at 2000 rpm, and then spin-cast the PDMS onto the molds at 266 rpm for 10 seconds then 1750-1850 rpm for 75 seconds. After allowing flow layers to rest on a flat surface for 10 minutes at room temperature, we baked them for 40 minutes at 80°C. Using a stereoscope, we then aligned the control layers to flow layers on master molds. The assemblies were baked for and additional 40 minutes at 80°C and then cut from the molds with a scalpel. Flow inlets were created with the same catheter hole punch.

#### Preparation of poly-L-lysine coated slides

PDMS devices do not adhere well to untreated glass. To functionalize glass surfaces to promote device adhesion and rounding of printed DNA droplets, we coated glass surfaces with poly-L-lysine. First, we washed 75 x 25 mm glass slides (Corning 2947) with Milli-Q H_2_O followed by drying. Next, we diluted Poly-L-lysine 0.1% (Electron Microscopy Sciences, 19320-A) 10-fold into Milli-Q H_2_O, added this solution to the glass slides, and allowed to shake for 1 hour at room temperature. Finally, we rinsed off the poly-L-lysine solution with Milli-Q H_2_O, baked them in a convection oven for 1 hour at 60°C, and then stored them in the dark at room temperature until use.

#### Plasmid printing and device alignment

To prepare plasmid solutions for spotting onto slides, we mixed 4 uL of each plasmid with 36 uL of filtered print buffer solution (0.5 g/L BSA, 100 mM NaCl, 12 g/L Trehalose) in a 384 well plate (Thermo Scientific, AB-1055); we stored these solutions at −20°C when not in use. Prior to printing, we defrosted the plate, centrifuged at 4°C (2000g for 10 minutes), and then carefully removed the foil to prevent cross-contamination. We printed plasmids onto the poly-L-lysine-coated glass slides using a sciFLEXARRAYER S3 fitted with a PDC70 nozzle with Type coating (Scienion). After printing, we aligned PDMS devices onto the printed arrays with each plasmid spot encapsulated by a different DNA compartment. Prior to use, we bonded the glass slides and PDMS devices together by baking for 3 hours at 95°C on a hotplate (Torrey Pines Scientific).

#### Microscopy instrumentation

We imaged all microfluidic devices using a Nikon Ti-S Microscope equipped with a motorized XY stage (Applied Scientific Instrumentation, MS-2000 XYZ stage), cMOS camera (Oxford Instruments, Andor Zyla 4.2 CMOS), and solid-state light source (Lumencor, Sola SE Light Engine), and automated filter turret with an eGFP filter set (Chroma Technology Corp., part no. 49002). We performed imaging with a 4X objective (CFI Plan Apochromat l 4X NA 0.20, Nikon) at 2×2 binning (1024×1024 pixels) with 500 ms exposure times. Throughout assays, we measured apparatus temperatures using a thermistor sensor (Thorlabs Inc., catalog no. TSP01, data logger; catalog no. TSP-TH NTC, external thermistor).

#### Conjugation of a SNAP substrate benzyl group (BG) to BSA

To covalently couple SNAP-substrate (benzyl guanine, BG) to BSA via N-hydroxysuccinimide (NHS) coupling, we dissolved NHS-GLA-BG (2 mg, S9151S, NEB) in 100 µL DMF, slowly added 20 µL of this solution to 1000 µL of 5 g/L buffer-exchanged BSA in 10 mM sodium borate (pH 8.0), and stirred wrapped in tin foil overnight. The BG-BSA was divided into 50 µL aliquots, flash frozen in liquid nitrogen, and stored at −80°C.

#### Surface patterning for construct immobilization

We connected Tygon tubing to a syringe at one end using a 23-gauge Luer connector (McMaster-Carr, 75165A684) and attached to a blunt steel pin at the other end (0.013 in ID x 0.025 in OD x 0.5 in length, New England Small Tube Corporation, NE-1310-02). With the reagents loaded into the syringe, we inserted the pin into the corresponding flow inlet on the microfluidic device. We then moved the other end of the tubing from the syringe to a custom pneumatic manifold, where fluid flow rates are controlled by pressure^100^. To actuate control valves, we used a pneumatic control manifold^100^ and Python software package (https://github.com/FordyceLab/RunPack)^101^.

To prevent premature solubilization of the DNA spots by osmosis, we pressurized control lines for the button, sandwich, and neck valves (**Figure S1**) with 550mM NaCl in Milli-Q H_2_O and remaining control lines with Milli-Q H_2_O. We used pressures from 25-27 psi to control device pneumatic valves and 2.5-3 psi to introduce reagents.

After dead-end filling all control lines, we pressurized (closed) button and neck valves to protect a circular region on device surfaces from fluid and block flow to the DNA chambers. To reduce nonspecific binding, we then coated all remaining surfaces with ultrapure bovine serum albumin (BSA) (5 g/L, ThermoFisher Scientific, AM2616). After flowing the BSA through the inlet tree to waste for ∼5 minutes, we opened the inlet valve to passivate all channels for ∼5 minutes. We closed the outlet valve and continued to apply pressure for an additional 10 minutes to expel all air from the device flow layer, and flowed BSA over the chip for an additional 15 min. Next, we flushed the device with phosphate buffered saline (10X stock, Corning, 46-013-CM; diluted to 1X in Milli-Q H_2_O) for 15 minutes. We then opened button valves and introduced BG-BSA (prepared as described above) for 20 minutes to specifically pattern the region protected by this valve with BG-BSA, followed by another PBS wash for 10 minutes.

#### On-chip expression of double-tagged constructs

We solubilized printed DNA spots encoding double-tagged constructs and expressed protein using the PURExpress expression system (New England Biolabs Inc., catalog no. E6800L). To prepare the expression mixture, we mixed 60 µL of PURExpress Solution A with 45 µL of Solution B and allowed this mixture to incubate on ice for 30–45 minutes. Immediately before flowing the reaction mixture into the chip, we added 3 µL of 40 U/L RNasin ribonuclease inhibitor (Promega Corporation, catalog no. N2515), 3 µL of 100 mM DTT, and 39 µL of nuclease-free water, and mixed gently. With the neck valves closed, we flowed this expression mixture over the device for 15 minutes to ensure that PURExpress solution fully filled all channels. To introduce the PURExpress mixture into all expression chambers and solubilize the DNA, we closed the chip outlet valve, opened the neck valves, and increased the flow pressure from to 4.0-4.5 psi, dead-end filling the DNA chambers under pressure. We monitored this process with a stereo microscope until chambers were 90–95% full and then closed the button, neck, and sandwich valves, isolating adjacent protein chambers from one another and preventing premature binding of constructs in the case of any leakage between expression chambers within the flow path. We then placed the slide bearing the device onto a pre-warmed aluminum hot plate (Torrey Pines Scientific) and allowed constructs to express for 2 hours at 30°C. Following expression, we removed devices from heat and incubated them for 90 minutes at room temperature in the dark to allow for maturation of the eGFP fluorophore.

#### Surface immobilization of expressed constructs

After construct expression, we removed the device/slide assembly from the hot plate and mounted it on the automated Nikon Ti-S microscope (components described above). To bind expressed constructs to the BG-BSA-patterned regions of the glass, we closed sandwich valves (to isolate adjacent reaction chambers from one another), opened button valves (to expose binding surfaces), and then opened the neck valve, allowing diffusion of expressed constructs into the reaction chambers. We allowed proteins to form covalent bonds between the SNAP tag and the BG-patterned surface for 30–60 min. During binding, we imaged chambers across the device in the eGFP channel to monitor binding, then closed button valves after observing minimal change in construct concentration.

#### Buffer exchange and thermolysin washing

Accurately measuring unfolding rates requires that chambers be free of contaminating constructs from other chambers and cell-free expression components, which can nonspecifically bind to device or slide surfaces. After closing button valves to protect immobilized protein, we opened sandwich valves to enable flow throughout the device, washed DNA and reaction compartments with Tris buffer (100 mM Tris•HCl (pH 8.0), 150 mM NaCl, 1 mM DTT) for 1 hour, and then closed the neck valves to isolate reaction compartments from DNA compartments.

To dislodge any nonspecifically bound proteins from underneath the button valves, we opened and closed the button valves 10 times, flowed Tris buffer for 10 minutes, then repeated this process 2 more times. With the button and neck valves closed, we increased the control pressure to 28-30 psi to protect the construct bound underneath the button from premature exposure to thermolysin. We then flowed 0.1 g/L thermolysin (prepared as described above) onto the device for 10 minutes to remove any cleavable material outside of the button regions.

#### Imaging and quantification of expressed constructs

To quantify construct expression and proteolytic degradation, we imaged microfluidic devices using an eGFP filter set (Chroma Technology Corp., part no. 49002) with acquisition times of 500 ms per image. Using a Python software package (https://pypi.org/project/acqpack/)^102^, each device was imaged using 2×2 binning and typically required 49 tiled images with 10% overlap. To assign mutant identities to each chamber, we stitched the overlapped tiles, divided each stitched image into 24×46 sub-images (stamps) centered on and containing a single chamber (using the 4 corner chambers as reference positions), determined the identity of the mutant at each position using a reference list, and then grouped images by mutant to process timecourses. Prior to the start of each proteolysis assay, we quantified starting eGFP intensities by: *(1)* identifying the centroid positions of button valve regions within each stamp using a combination of a sparse grid search (to identify the circular region of maximal summed fluorescence intensity) and a dense local grid search (to refine this position), *(2)* quantifying the median intensity from all pixels within a fixed radius of 15 pixels (larger than the physical radius of the button), and then *(3)* subtracting the local background by calculating the median intensity of pixels in an annulus concentric with, abutting, and larger than the button bounding circle. For fitting photobleaching rates, intensities were not background-subtracted.

#### Measuring unfolding rates on-chip

To quantify rates of proteolytic degradation, we: *(1)* loaded 0.1 g/L thermolysin into each chamber, *(2)* closed sandwich valves, *(3)* opened the buttons (to expose surface-immobilized protein to thermolysin), and *(4)* started automated image acquisition with a Python software package (https://pypi.org/project/acqpack/)^102^. Images were collected from time point 0 to 10,000 s; images were acquired more frequently at the beginning of the assay (to capture fast cleavage for constructs such as AGLA) and less frequently at the end of the assay (to capture slow cleavage for constructs such as DGLP while limiting photobleaching). As low-expressing mutants yielded unfolding rates with higher variance, data were filtered to only report values for variants with >5 chambers with measured initial intensities > 300.

#### Testing statistical significance of measured effects

To assess the statistical significance of measured differences in unfolding rates, we performed bootstrapped hypothesis testing for each pairwise comparison between the set of measured unfolding rates for a given mutant *vs.* the WT construct. The null hypothesis for each test was that there was no difference in unfolding rates. To account for multiple comparisons, we applied a Bonferroni correction to adjust for the significance threshold by calculating a Bonferroni-corrected p-value threshold (α/N, where α = 0.05 and N was the total number of mutants tested). Mutants with significant *p*-values were classified as having a statistically different unfolding rate compared to the WT.

#### Estimation of time and cost for traditional vs. SPARKfold methods

In a traditional assay, transforming the DNA encoding the desired protein mutants into bacteria and plating the cells to allow colony growth requires about 1.5 days, followed by 2 days to grow the cell cultures. Lysing cells and purifying the target proteins require 1 additional day. Buffer exchanging to transfer the protein into a compatible buffer for downstream assays takes 0.5 days. Unfolding assays, which can involve approaches with plate reader and SDS-PAGE gel readouts, require about 2 days, totaling 7 days. Assuming that 5 mutants can be prepared and assayed in parallel, 500 mutants can be studied in 700 days. If the salary for one person is $62,500 per year (reflecting typical rates for graduate students and postdoctoral researchers), and there are 250 workdays per year, the labor cost for two people to complete 700 days of work each is $350,000. In contrast, SPARKfold can simultaneously express and purify 500 mutants on a microfluidic chip within just 1 day, followed by buffer washes and unfolding assays completed in the following day. Given the same salaries and workdays, the cost for 2 people to perform this 2-day assay is $1,000, a savings of >100 fold.

#### ecDHFR variant analyses

For ecDHFR mutations made in alpha helices, we determined the degree to which unfolding rates could be explained by predicted changes in helical propensity by plotting measured unfolding rates vs. predicted ΔΔG helical propensity values (calculated as Δ*G* for substituted residue -Δ*G* for native residue, using previously-reported amino acid helical propensities^76^) and performed a linear regression. The relative solvent accessibility (RSA) of residues in the ecDHFR protein was calculated using PyMOL’s get_sasa_relative function. This function computes the relative solvent accessible surface area for each residue, with values ranging from 0.0 (fully buried) to 1.0 (fully exposed). To assess ecDHFR conservation, we created a multiple sequence alignment (MSA) from PFAM (PF00186), used BioPython to align each sequence to the reference species and remove gaps, and then calculated a frequency matrix of amino acid occurrences across the positions. We visualized conservation by plotting the maximum frequency at each ecDHFR position and creating sequence logos using WebLogo^103^. For the Φ analysis, we calculated the free energy for partial unfolding (ΔΔ*G*_C-N_°) using median SPARKfold values for variant and WT ecDHFR, then determined Φ by using this measured ΔΔ*G*_C-N_° and previously published measurements of free energy for global unfolding (ΔΔ*G*_U-N_°)^43–47,68^, as described by Park^43^.

