## Supplementary Information for "Quantifying protein unfolding kinetics with a high-throughput microfluidic platform"

Atsavapranee, B.[1*], Sunden, F.[2*], Herschlag, D.[2], & Fordyce, P.M.[1,3,4,5]
[1]Department of Bioengineering, Stanford University, Stanford, CA 94305
[2]Department of Biochemistry, Stanford University, Stanford, CA 94305
[3]Department of Genetics, Stanford University, Stanford, CA 94305
[4]Sarafan ChEM-H Institute, Stanford University, Stanford, CA 94305
[5]Chan Zuckerberg Biohub, Stanford University, Stanford, CA 94305
*These authors contributed equally

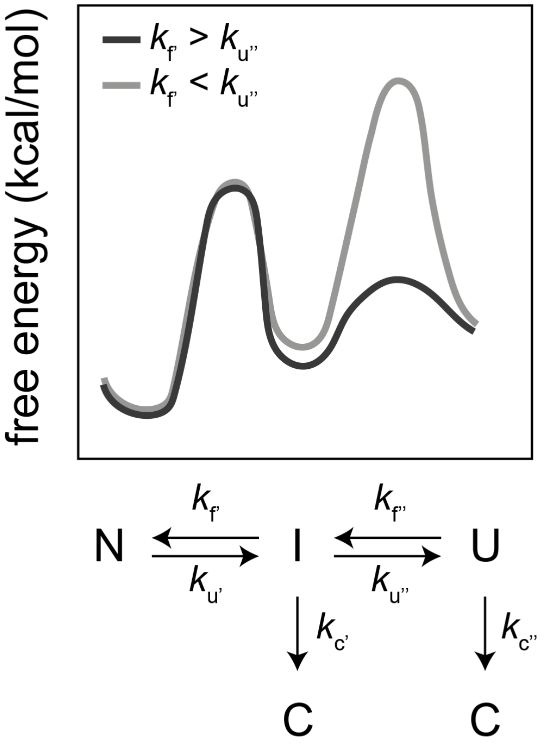


**Figure S1 (Related to Figure 1). Free energy diagram (top) and reaction scheme (bottom) for an unfolding pathway with a native state (N), an unfolded state (U), and a single unfolding intermediate (I).** Even for non-two-state proteins with unfolding intermediates, monitoring proteolysis under conditions where any unfolded protein is immediately cleaved can reveal the rate of the first unfolding step (provided that intermediates are susceptible to proteolytic cleavage).

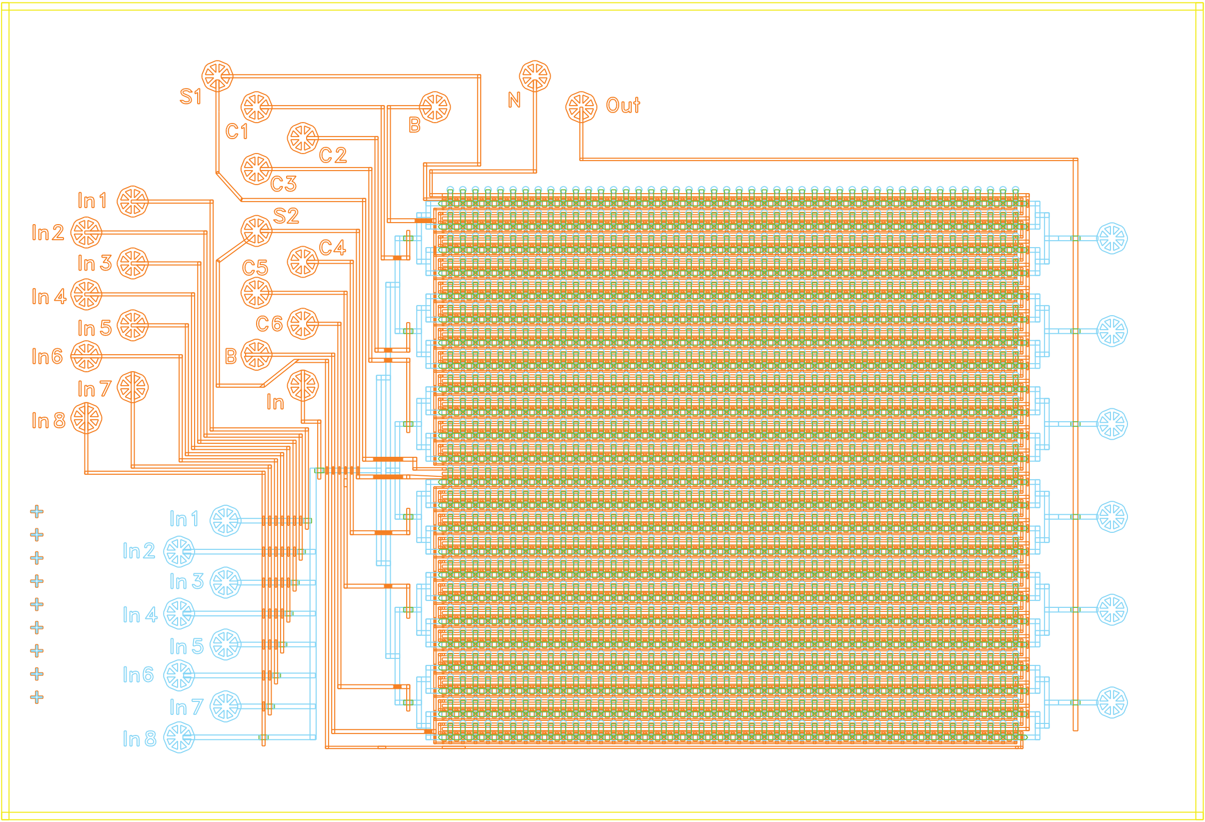


**Figure S2 (Related to Figure 1).** Architecture of SPARKfold microfluidic device. Detailed diagram of SPARKfold microfluidic device with valve inlets and outlets showing reagent flow channels (blue) and pneumatic valve channels (orange) that control flow of reagents in device.

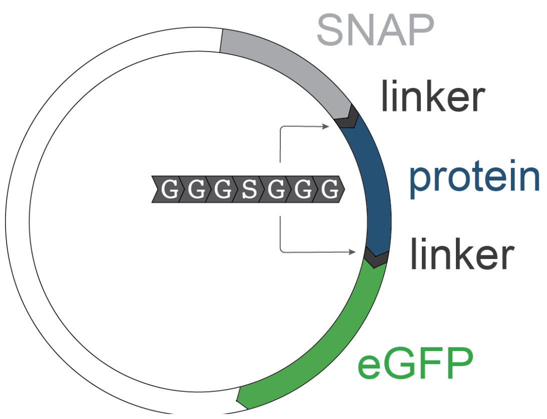


**Figure S3 (Related to Figure 1).** Schematic of plasmids used for expression of variants with a N-terminal SNAP tag and a C-terminal eGFP tag connected by flexible glycine/serine linkers.

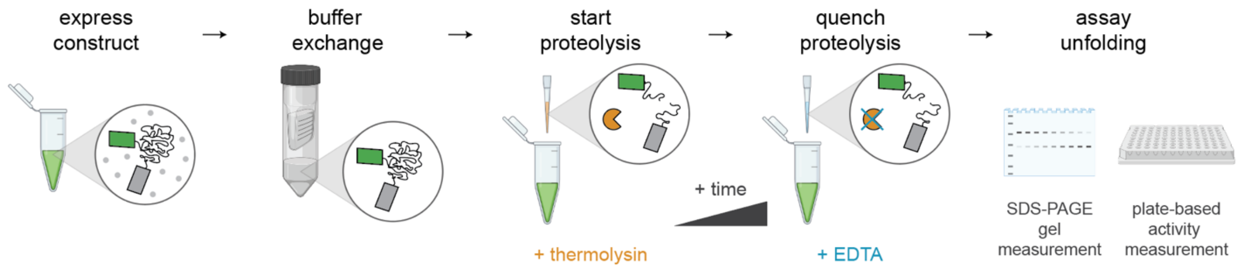


**Figure S4 (Related to Figure 3).** Cartoon schematic showing traditional native proteolysis assay workflow: (1) protein constructs are expressed and buffer exchanged in solution, (2) thermolysin protease (orange) is added to start proteolysis, (3) ethylenediaminetetraacetic acid (EDTA) (blue) is added to quench proteolysis at varying times, and then (4) samples are analyzed with sodium dodecyl sulfate-polyacrylamide gel electrophoresis (SDS-PAGE) or plate-based activity measurements to quantify the amount of protein cleaved as a function of time.

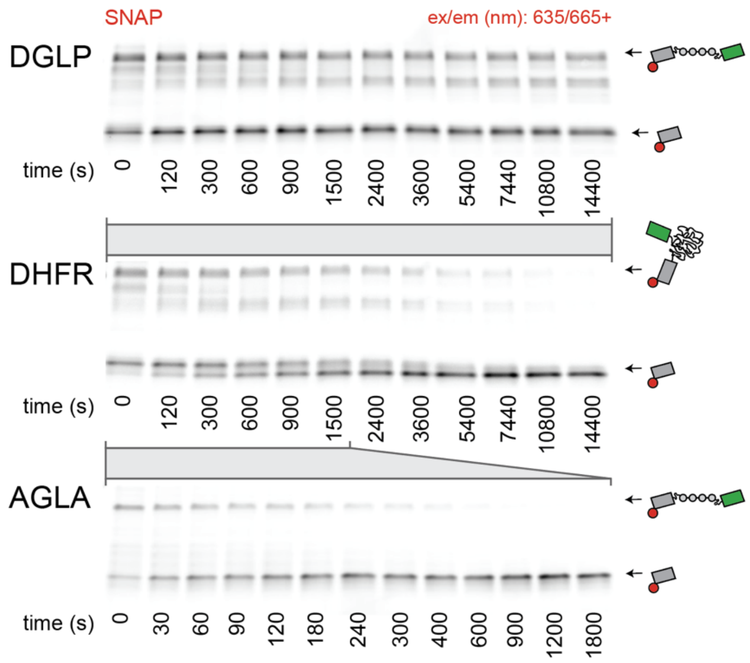


**Figure S5 (Related to Figure 3).** Fluorescence images of full SDS-PAGE gels showing amount of full-length protein constructs remaining after native proteolysis in 0.1 g/L thermolysin at various time points. Protein constructs contain an N-terminal SNAP tag (grey rectangle) labeled with an SNAP Surface-649 fluorophore (red circle) and a C-terminal monomeric eGFP fluorescent protein (green rectangle) surrounding either a negative control sequence designed to be minimally cleaved by thermolysin (DGLP, top), WT ecDHFR (middle), or a positive control sequence comprised of a preferred thermolysin cleavage site (AGLA, bottom). Images were collected at an excitation wavelength of 635nm and emission wavelengths above 665nm to quantify SNAP-labeled protein construct fragments with increasing time exposed to thermolysin; arrows and associated cartoons indicate full-length protein constructs and free dye-labeled SNAP tags.

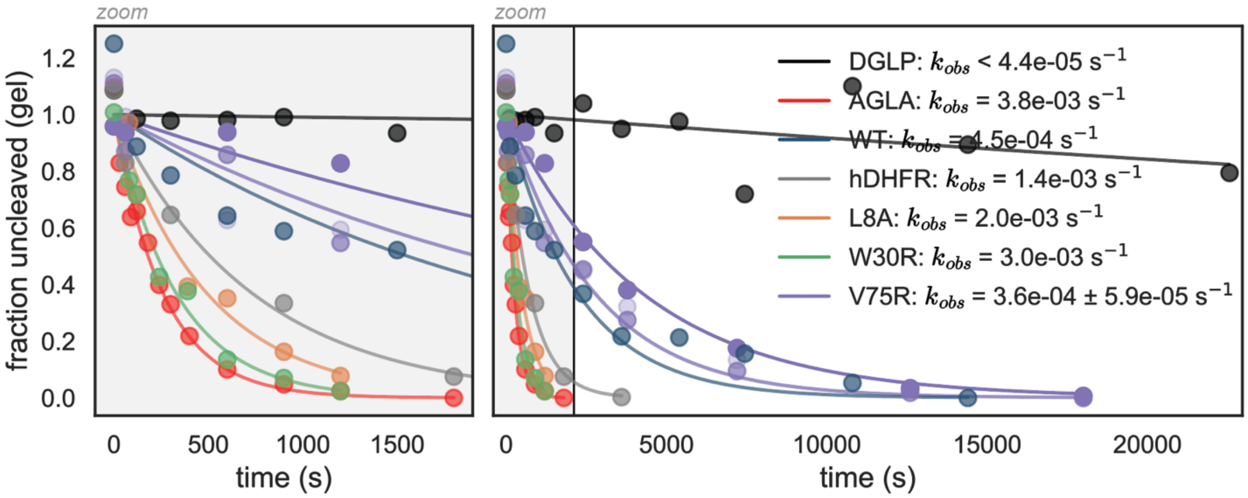


**Figure S6 (Related to Figure 3).** Fraction of uncleaved protein remaining as a function of time after traditional native proteolysis using an SDS-PAGE gel electrophoresis readout for a subset of control and DHFR variant constructs. The lefthand plot shows the first 2000 seconds of the timecourse and the righthand plot shows the full time course. Markers represent normalized intensities of the full-length protein band at a given time point and lines indicate single exponential decay fits to plotted data. The measured cleavage rate for DGLP represents a lower limit. For ecDHFR V75R, data is shown for experiments carried out at 0.06 g/L (light purple), 0.3 g/L (medium purple), and 1g/L (dark purple), and the legend reports the median and standard deviation of the unfolding rates across these thermolysin concentrations. For all other variants, measurements were performed with 0.1 g/L thermolysin (matching the concentration used in SPARKfold assays).

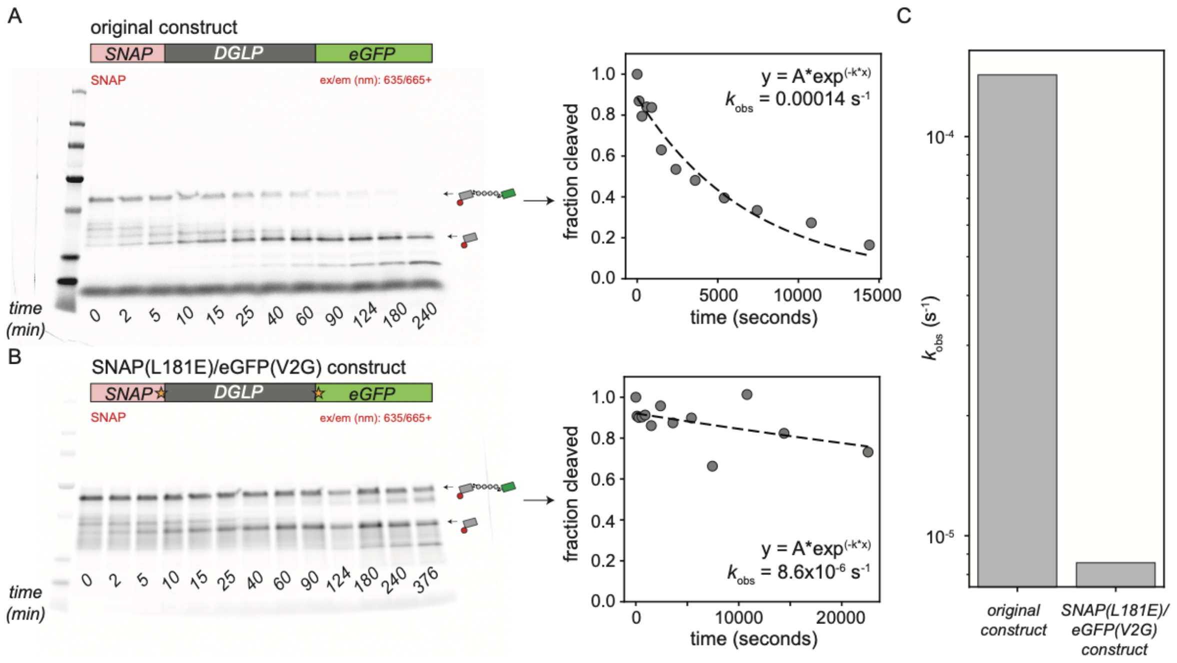


**Figure S7 (Related to Figure 3).** Mutations designed to remove potential thermolysin cleavage sites from SNAP and eGFP protein tags (SNAP L181E and eGFP V2G, respectively) increase resistance to proteolysis. **(A)** Fluorescence image after gel electrophoresis (left) and scatter plot quantifying fraction cleaved as a function of proteolysis time (right) for the original construct containing the DGLP negative control sequence flanked by N-terminal SNAP and C-terminal eGFP tags. Cartoon schematics indicate expected band identities, scatter plot markers represent normalized intensities of the full-length band, and dashed line shows a single exponential fit to the data. **(B)** Fluorescence image after gel electrophoresis (left) and scatter plot quantifying fraction cleaved as a function of proteolysis time (right) for the SNAP L181E/eGFP V2G construct. Cartoon schematics indicate expected band identities, scatter plot markers represent normalized intensities of the full-length band, and dashed line shows a single exponential fit to the data. **(C)** Bar graph showing fitted time constants for the original and mutant constructs.

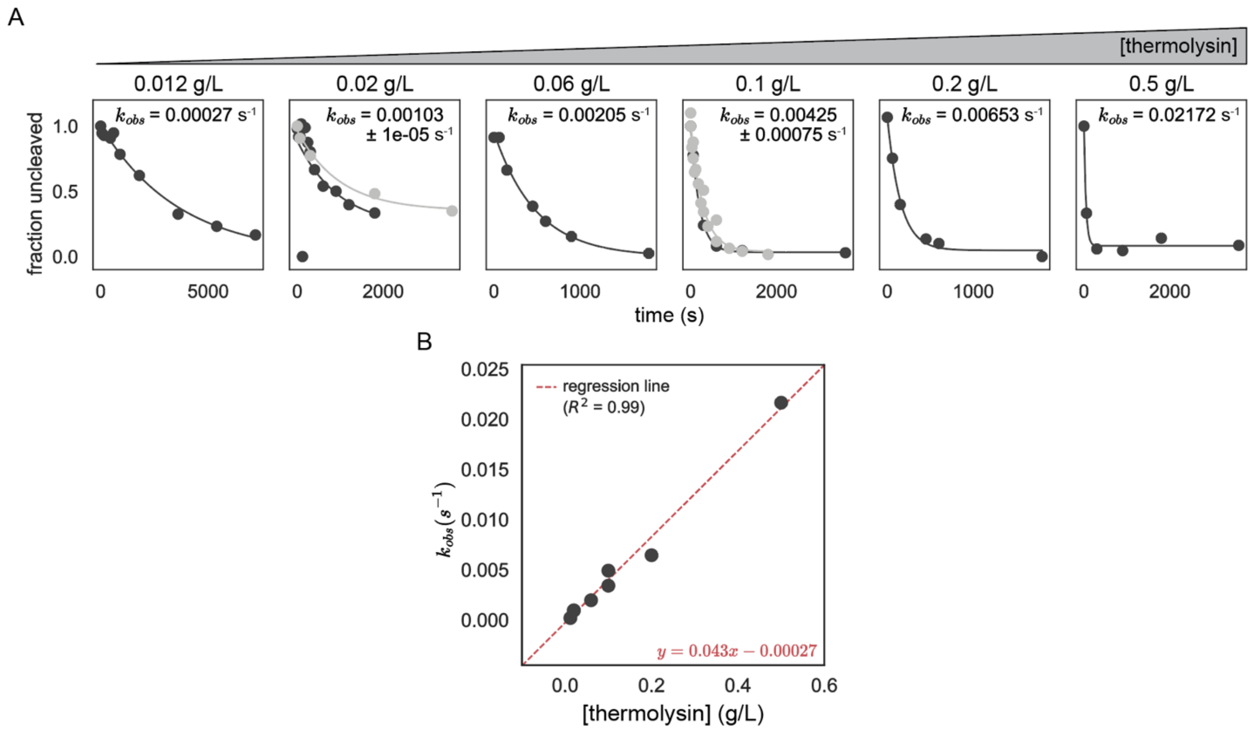


**Figure S8 (Related to Figure 3).** Cleavage rates of a linear (always unfolded) thermolysin-preferred substrate (AGLA, positive control) vary linearly with thermolysin concentration as expected for first order kinetics. **(A)** Fraction uncleaved (*y* axis) *vs.* proteolysis time (*x* axis) as a function of increasing thermolysin concentrations from 0.012 g/L (left) to 0.5 g/L. Black and grey markers indicate results from replicate experiments; black and gray lines indicate single exponential decay fits to data. Annotated cleavage rates are calculated as the median and standard deviation of the cleavage rates across replicates. **(B)** Fitted cleavage rate as a function of thermolysin concentration. Red dashed line indicates linear fit to plotted data ($R^2$ = 0.99).

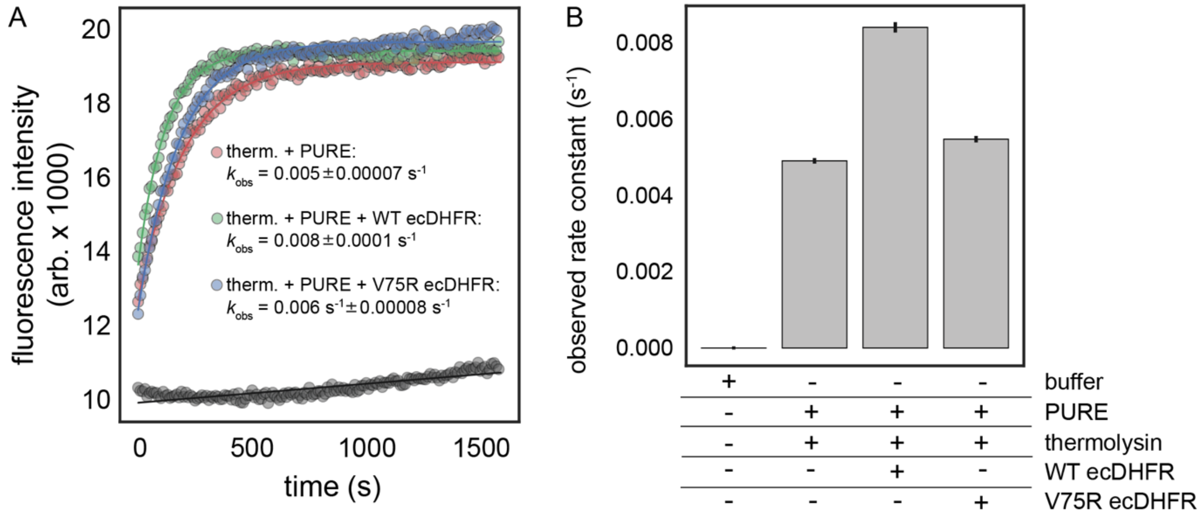


**Figure S9 (Related to Figure 3). Thermolysin activity in multi-well plate assays is not inhibited by components of the cell-free protein synthesis mix or the presence of expressed ecDHFR constructs.** **(A)** Measured fluorescence intensity as a function of time for buffer alone (grey markers), purified thermolysin with cell-free protein expression reagents without plasmid DNA (red markers), with plasmid DNA encoding WT ecDHFR (green), and with plasmid DNA encoding a thermodynamically destabilized ecDHFR variant (V75R). Progress curves were fit (solid lines) to a single exponential function with an offset: $y = A(1 - e^{-k_{obs}t}) + y_0$. **(B)** Bar chart showing fitted rate constant ($k_{obs}$) for each condition.

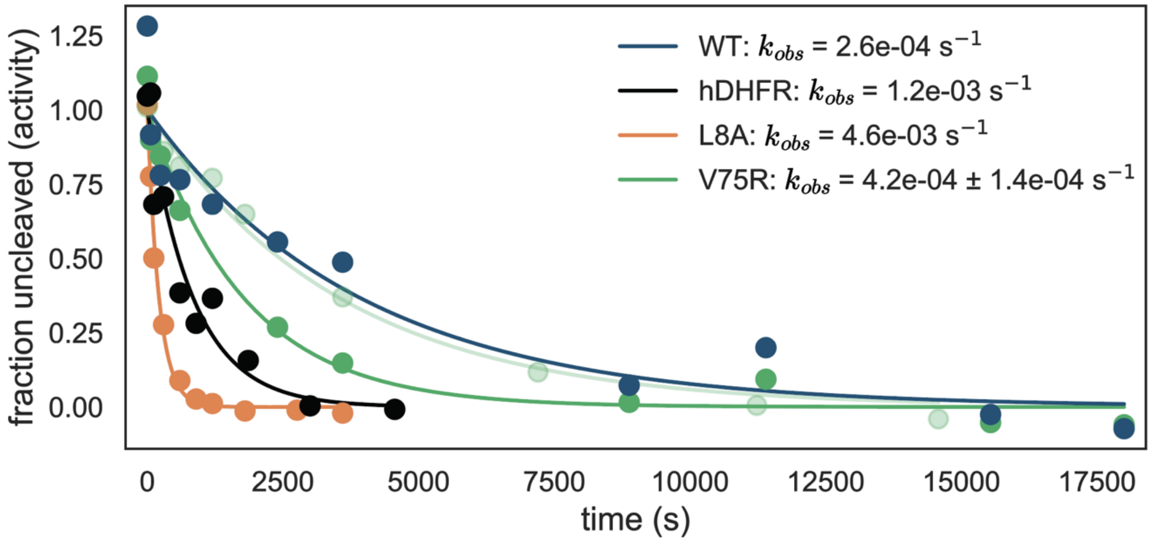


**Figure S10 (Related to Figure 3).** Quantifying fraction uncleaved via plate-based activity measurements following native proteolysis for SNAP- and eGFP-tagged WT ecDHFR, WT hDHFR, L8A ecDHFR, and V75R ecDHFR constructs. The activity at each time point was normalized to the initial activity to determine the fraction of uncleaved enzyme (*y* axis) after varying durations of proteolysis at 0.1 g/L thermolysin. Lines indicate single exponential decay fits to plotted data. For V75R, data is shown for replicate experiments (light and dark green). Annotated unfolding rates are calculated as the median and standard deviation of the cleavage rates across replicates.

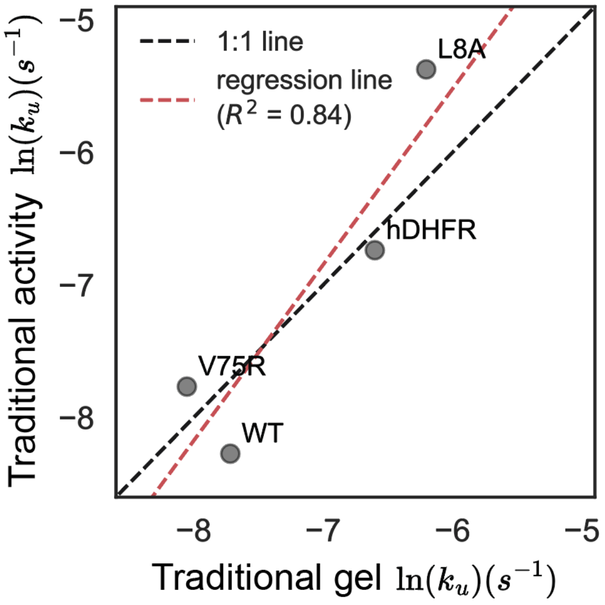


**Figure S11 (Related to Figure 3).** Comparison between unfolding rates determined from traditional native proteolysis followed by plate-based activity assays (*y* axis) *vs.* native gel electrophoresis (*x* axis) for WT *E. coli* DHFR (WT), human DHFR (hDHFR), and the V75R and L8A mutants of *E. coli* DHFR. Red dashed line indicates linear fit to plotted data ($R^2$ = 0.84), and black dotted line indicates the 1:1 relationship.

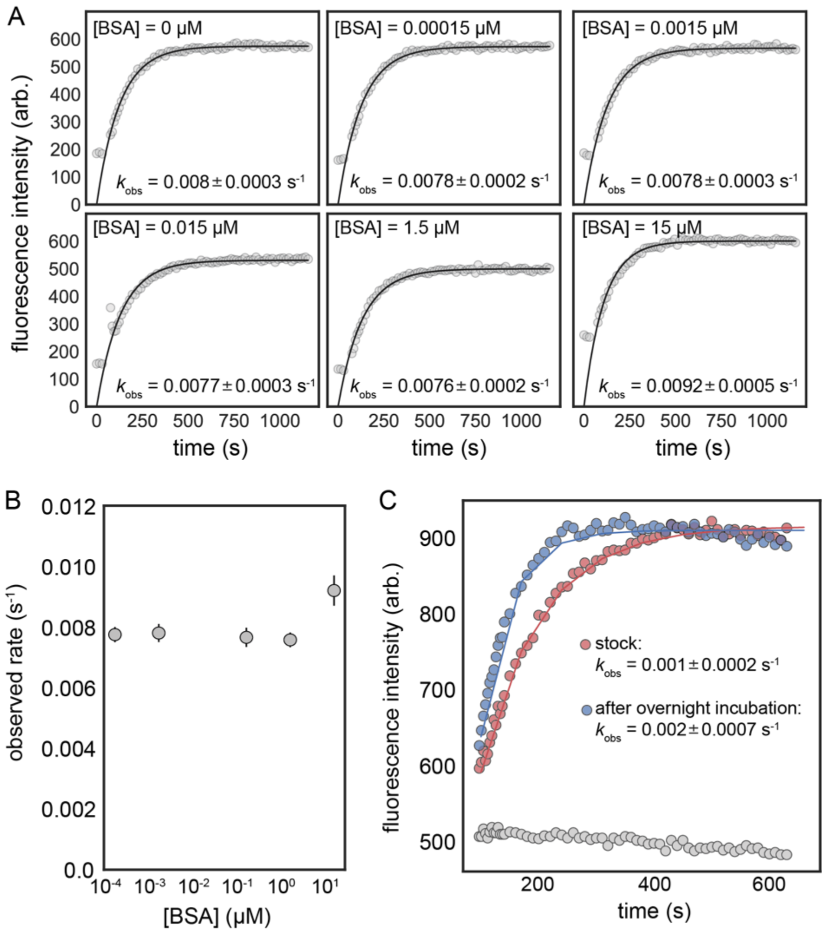


**Figure S12 (Related to Figure 4). Thermolysin activity is not inhibited by BSA or overnight incubation in Tygon tubing. (A)** Scatter plots showing measured fluorescence intensity over time for 0.1 mg/mL thermolysin in the presence of increasing concentrations of BSA. Progress curves were fit (solid lines) to a single exponential function with an offset: $y = A(1 - e^{-k_{obs}t}) + y_0$; fit parameters are indicated on each panel. **(B)** Observed rate as a function of BSA concentration. **(C)** Scatter plot showing measured fluorescence intensity over time for buffer alone (grey markers), 0.1 mg/mL thermolysin directly after thawing (‘stock’, red markers), and 0.1 mg/mL thermolysin after an overnight incubation in Tygon tubing (blue markers), simulating conditions of an on-chip SPARKfold assay.

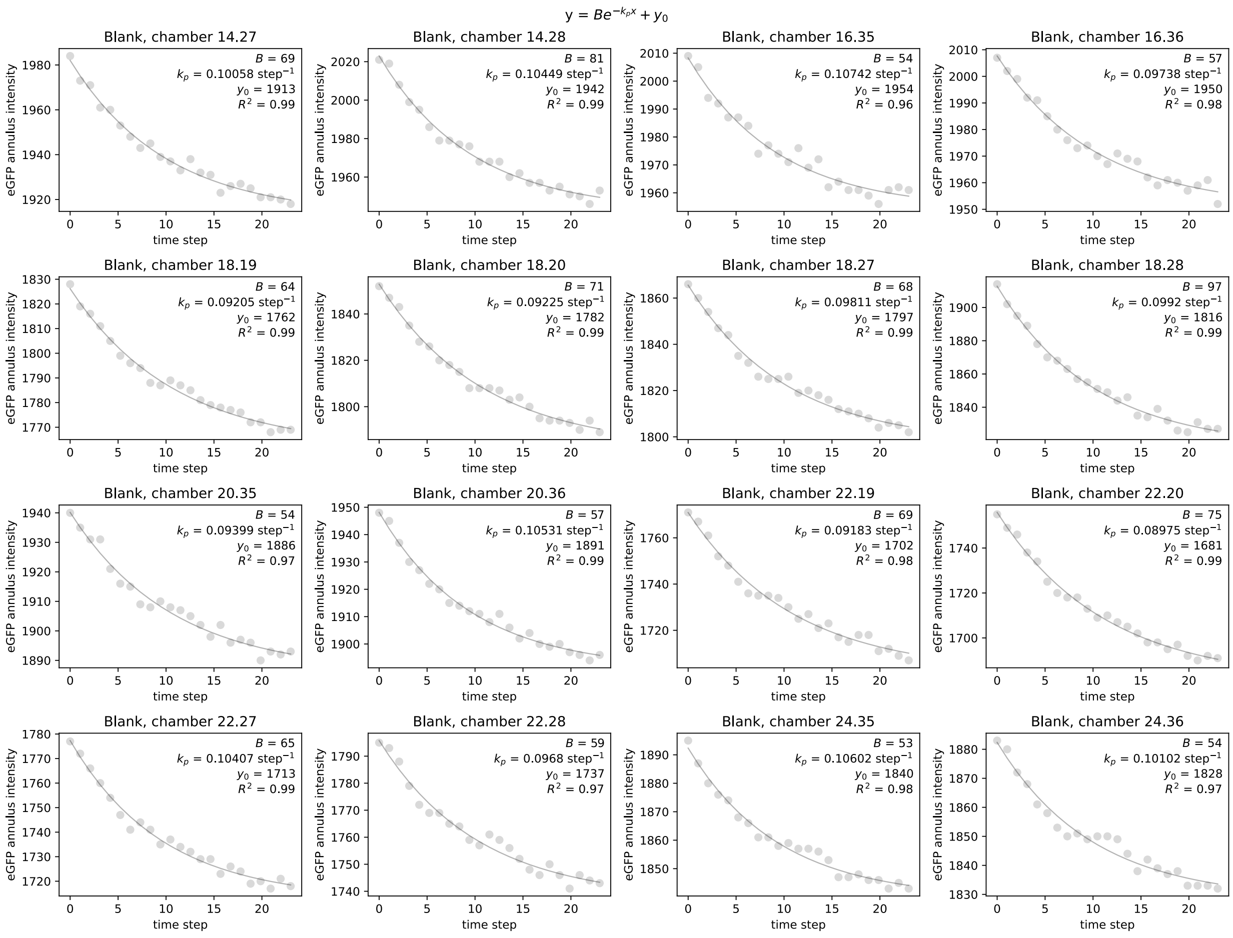


**Figure S13 (Related to Figure 4).** Example per-chamber eGFP fluorescence intensity decay curves for 16 ‘blank’ chambers from a single experiment; these data (unlike data for protein-containing chambers) reflect intensities with no background subtraction. Grey markers indicate measured eGFP intensities (arbitrary units) at each time point. Grey line indicates single exponential decay fit returning the annotated fit parameters for each chamber ($k_p$ = photobleaching rate).

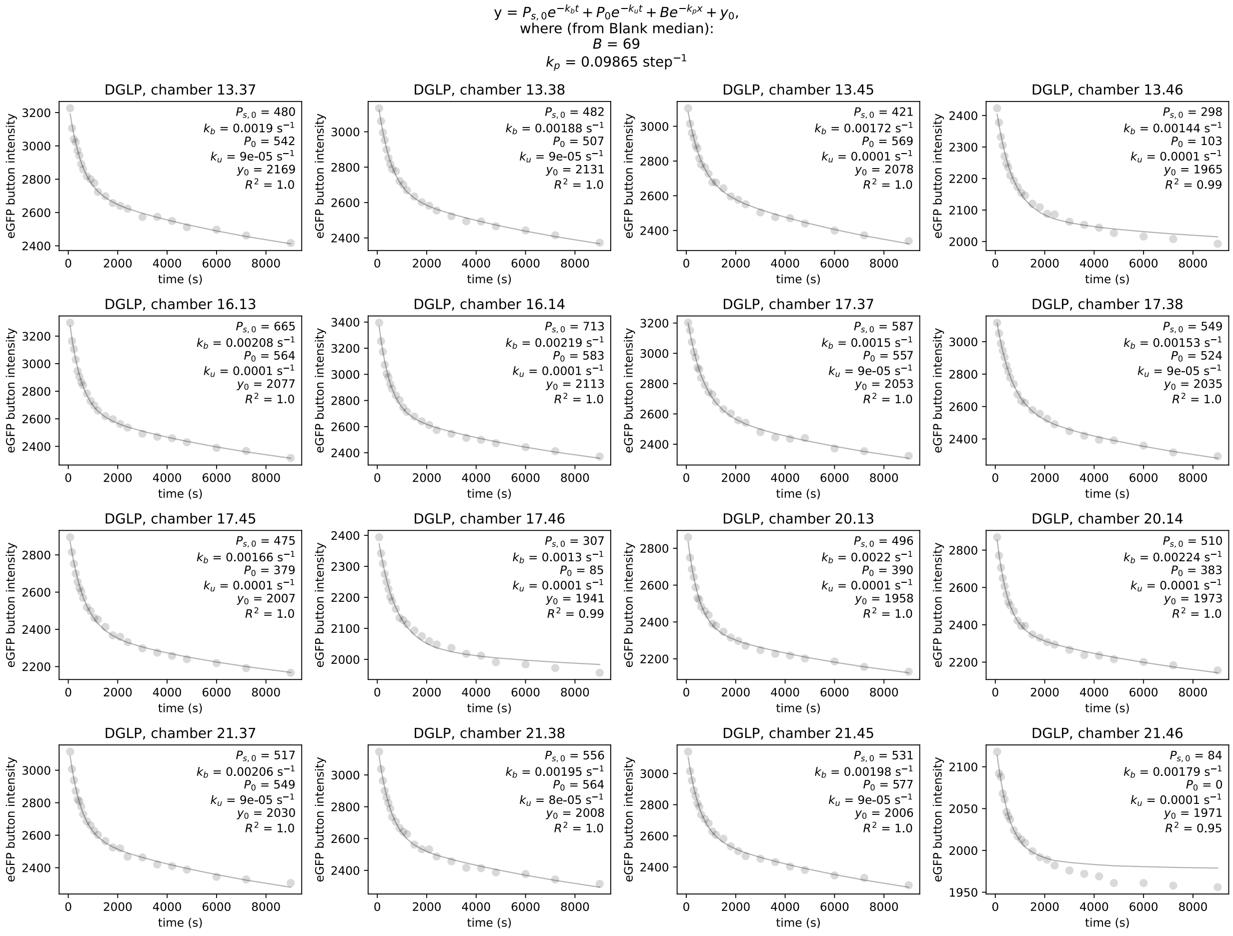


**Figure S14 (Related to Figure 4).** Example per-chamber eGFP fluorescence intensity decay curves for 16 chambers containing a linear non-thermolysin-preferred sequence (DGLP) within a single experiment. Grey markers indicate measured eGFP intensities (arbitrary units) at each time point. Grey line indicates fit to the function shown at top with the photobleaching rate constrained to the value obtained from blank chambers ($k_{\mathrm{b}}$ = rate of slow dissociation of weakly-bound population from surface, $k_{\mathrm{u}}$ = measured unfolding rate).

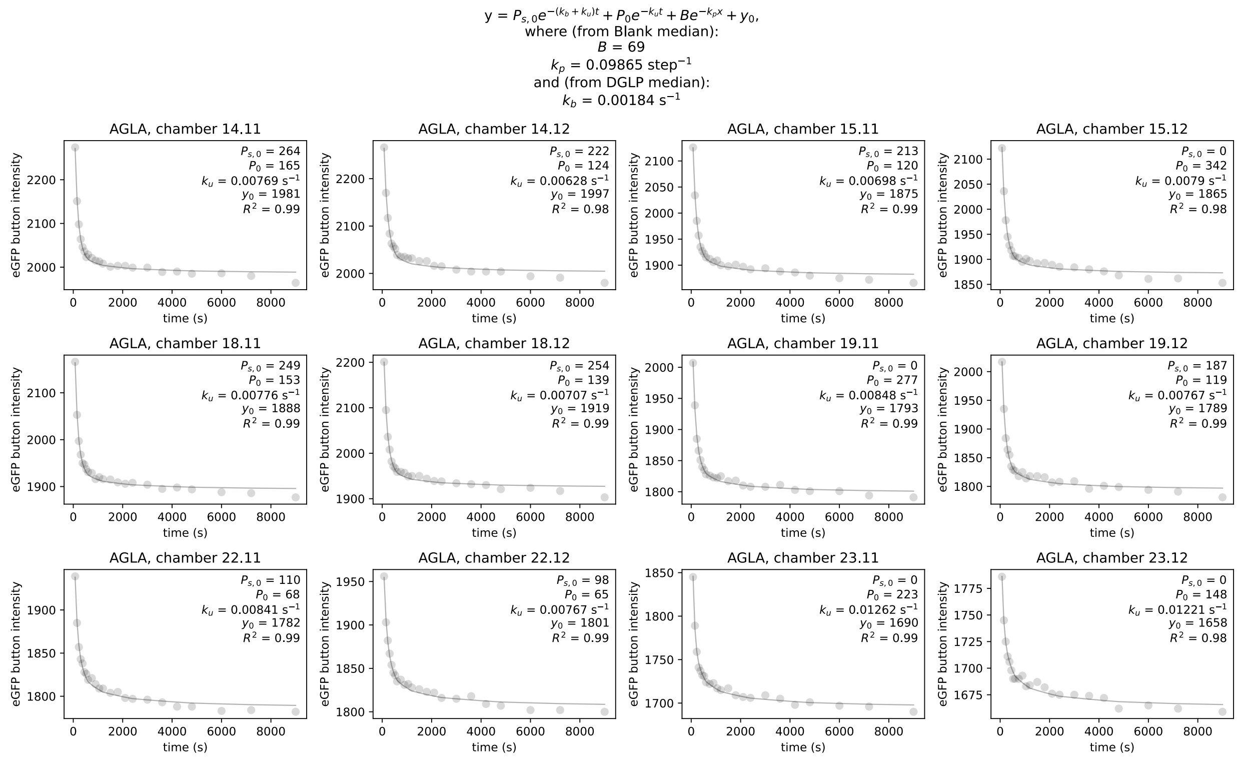


**Figure S15 (Related to Figure 4).** Example per-chamber eGFP fluorescence intensity decay curves for 12 chambers containing a linear thermolysin-preferred sequence (AGLA) within a single experiment. Grey markers indicate measured eGFP intensities (arbitrary units) at each time point. Grey line indicates fit to the function shown at top with the photobleaching rate constrained to the value obtained from blank chambers and the slow dissociation rate constrained to the value obtained from negative control (DGLP)-containing chambers ($k_u$ = rate constant for cleavage).

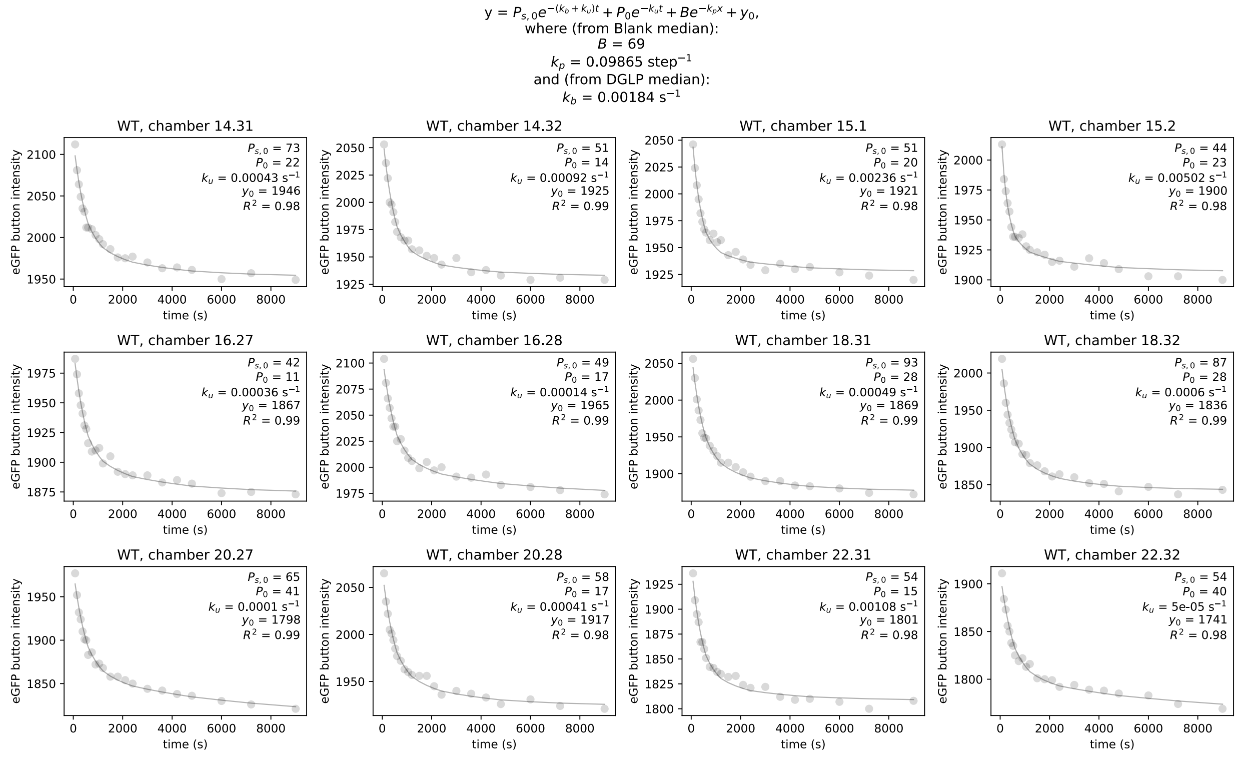


**Figure S16 (Related to Figure 4).** Example per-chamber eGFP fluorescence intensity decay curves for 12 chambers containing WT ecDHFR within a single experiment. Grey markers indicate measured eGFP intensities (arbitrary units) at each time point. Grey line indicates fit to the function shown at top with the photobleaching rate constrained to the value obtained from blank chambers and the slow dissociation rate constrained to the value obtained from negative control (DGLP)-containing chambers ($k_u$ = rate of unfolding).

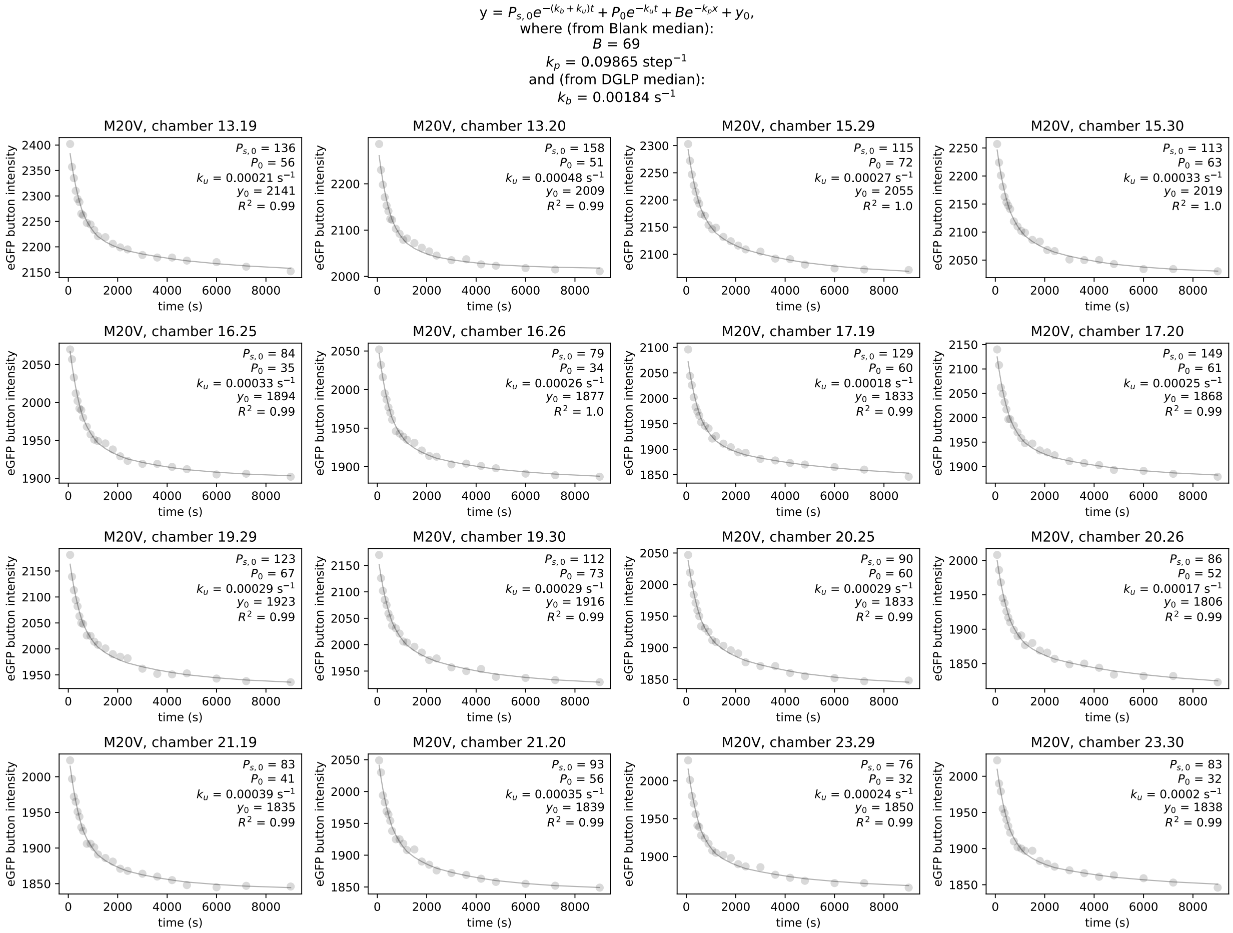


**Figure S17 (Related to Figure 4).** Example per-chamber eGFP fluorescence intensity decay curves for 12 chambers containing M20V ecDHFR within a single experiment. Grey markers indicate measured eGFP intensities (arbitrary units) at each time point. Grey line indicates fit to the function shown at top with the photobleaching rate constrained to the value obtained from blank chambers and the slow dissociation rate constrained to the value obtained from negative control (DGLP)-containing chambers ($k_u$ = rate of unfolding).

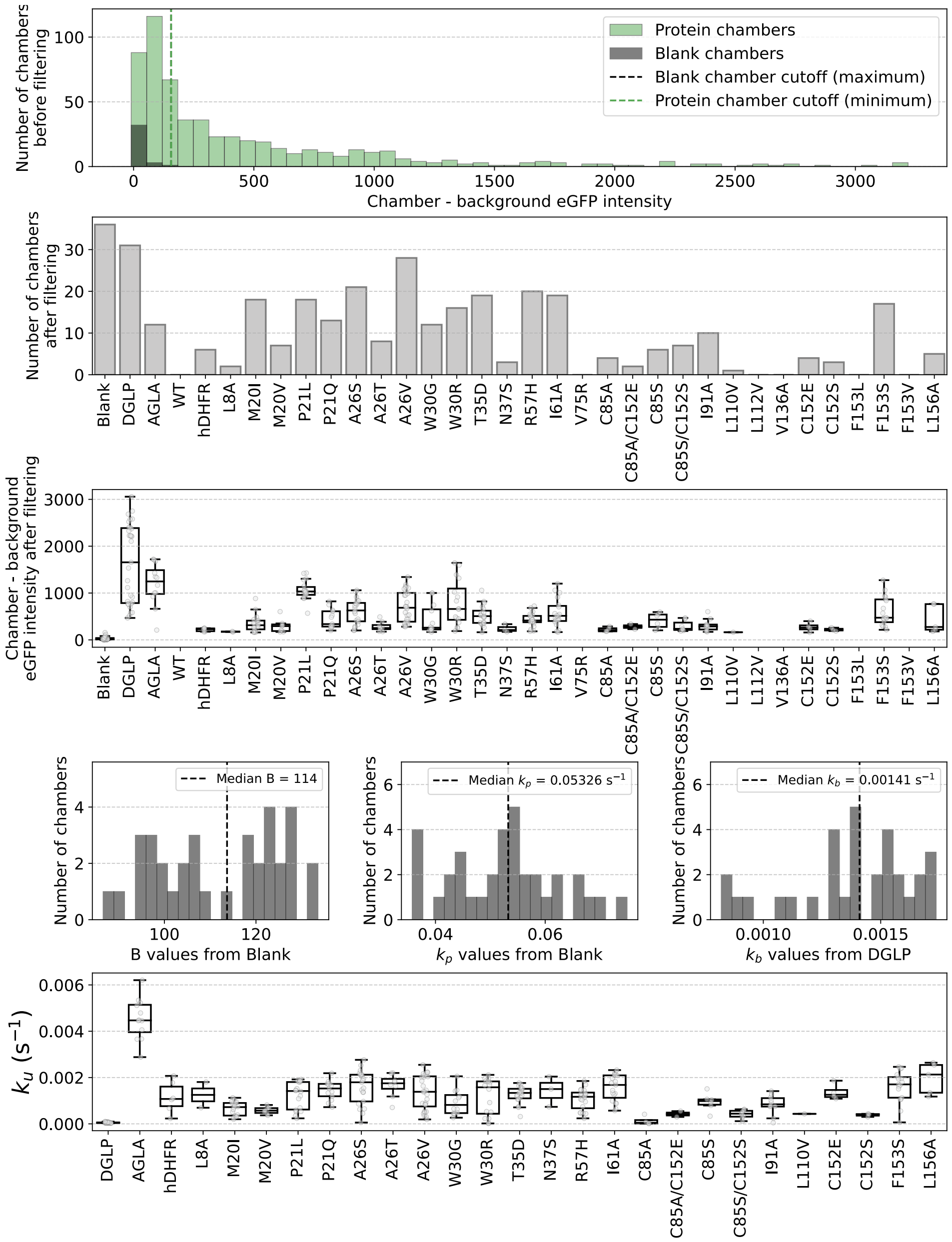


**Figure S18 (Related to Figure 4).** Example summary report for a single experiment showing: (1) histograms of measured eGFP intensities for plasmid-containing (green) and empty (grey) chambers and minimum threshold used to identify chambers with expressed protein (green dashed line), (2) number of chambers with intensities above the minimum threshold for each variant construct, (3) box and whisker plots showing the distribution of background-subtracted expression values for all chambers by variant, where the box spans the interquartile range (IQR), the line inside the box represents the median, whiskers extend to 1.5xIQR beyond the box, and points outside this range are shown as outliers, (4) histograms showing distributions of absolute offsets (left) and photobleaching rates (middle) from blank chambers and slow dissociation rates from negative control (DGLP)-containing chambers (right), and (5) box and whisker plots showing the distribution of fitted unfolding rates for all chambers by variant, where the box spans the IQR, the line inside the box represents the median, whiskers extend to 1.5xIQR beyond the box, and points outside this range are shown as outliers.

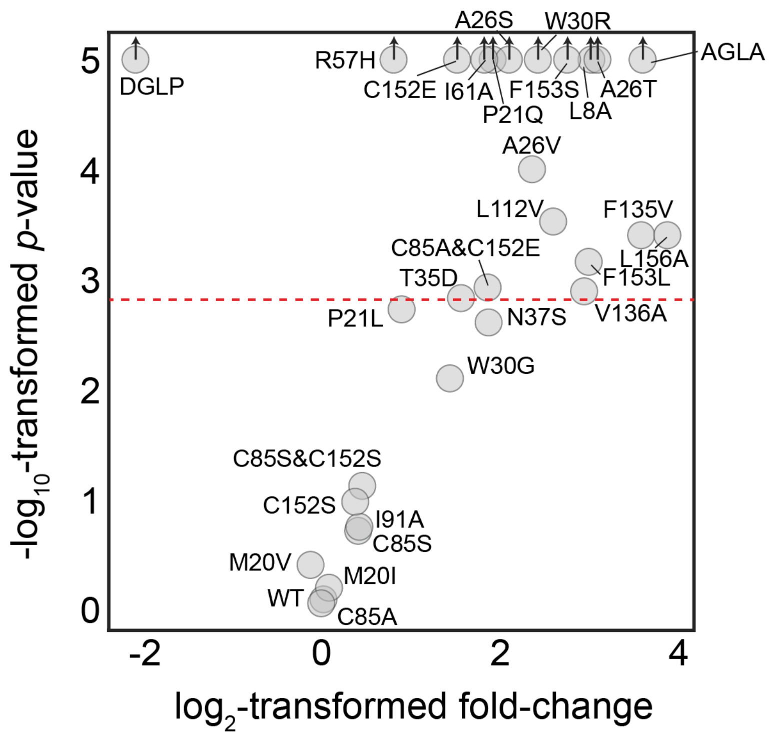


**Figure S19 (Related to Figure 4). Volcano plot of measured unfolding rates**. Scatter plot showing bootstrapped log10-transformed *p*-values vs. log2-trasformed fold-change in unfolding rates relative to WT ecDHFR. Red dashed line indicates Bonferroni-corrected significance. Mutants with $p = 10^{-5}$ represent upper limits of significance (indicated by an upward-facing arrow).

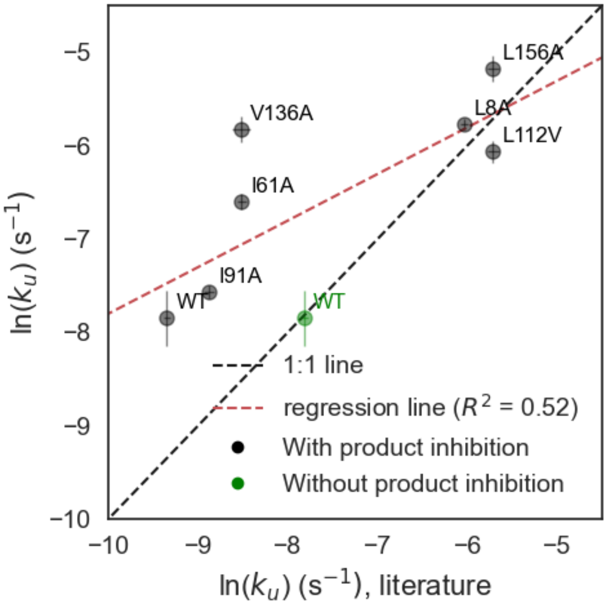


**Figure S20 (Related to Figure 4). Comparison between SPARKfold and previously published ecDHFR variant unfolding rates.** Data relates the natural logarithm of the median unfolding rate determined by SPARKfold with error bars indicating the standard error (*y* axis) and the natural logarithm of the published unfolding rate (x axis) for initial experiments with product inhibition(Kasper, Liu, et al., 2014) (black) and follow-up experiments without product inhibition(Kasper, Andrews, et al., 2014) (green). Red dashed line indicates linear fit to plotted data with product inhibition ($R^2$ = 0.51), and black dotted line indicates the 1:1 relationship.

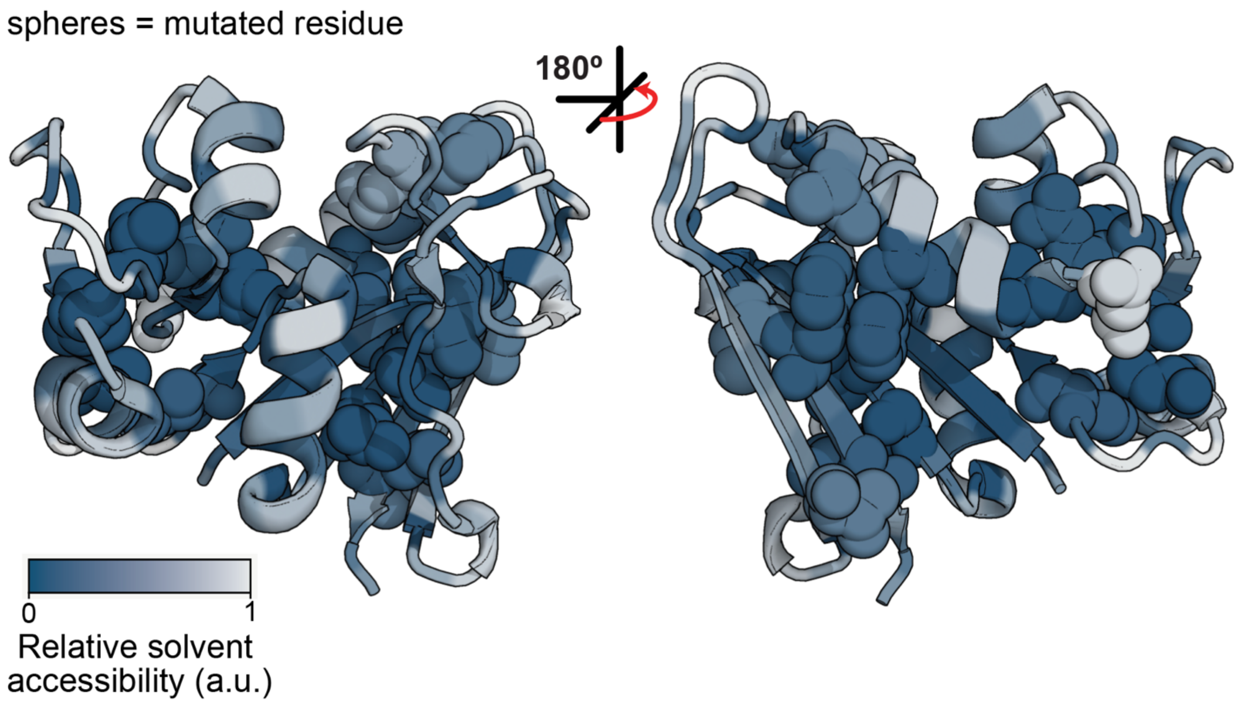


**Figure S21 (Related to Figure 5).** Relative solvent accessibility of residues on the ecDHFR structure. Residues mutated in this study are shown as spheres. All ecDHFR residues are colored on a scale from low (dark blue) to high (light gray) relative solvent accessibility.

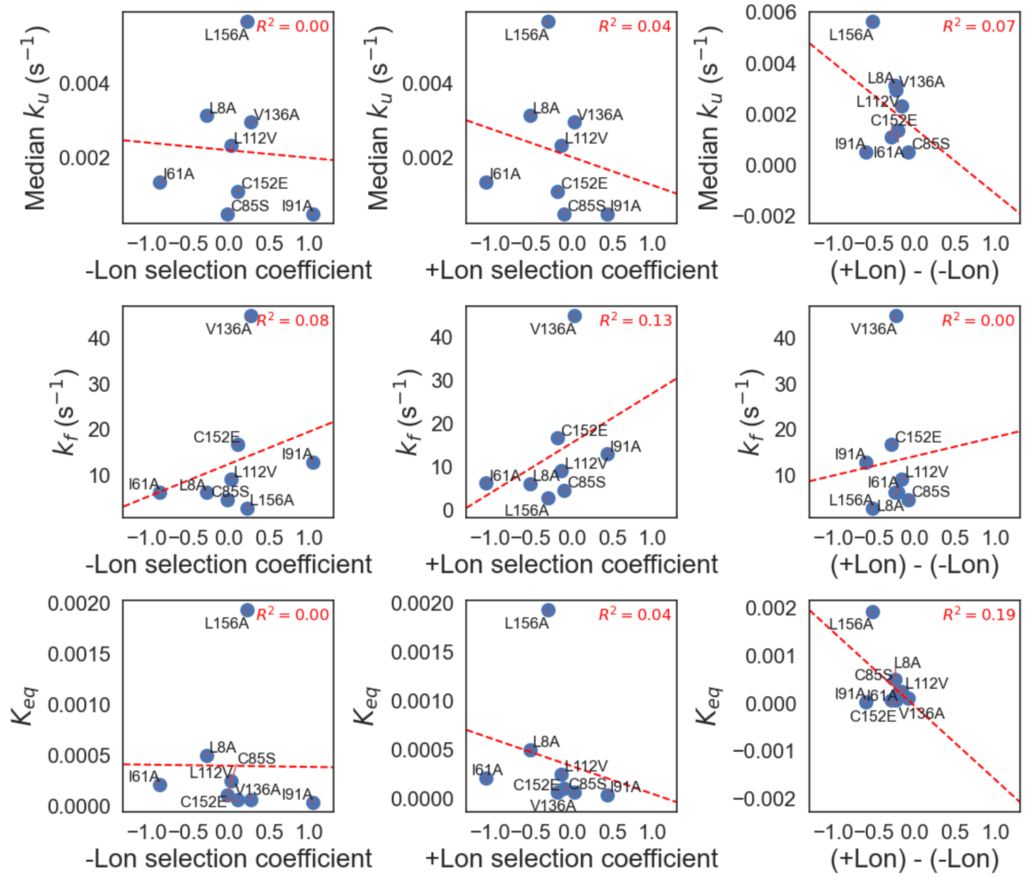


**Figure S22.** Comparison between ecDHFR kinetic and thermodynamic stability parameters and Lon selection coefficients. For ecDHFR variants with published equilibrium constant ($K_{eq}$) values, scatter plots display the measured median unfolding rate ($k_u$) and previously published $K_{eq}$ values(Garvey & Matthews, 1989; Iwakura et al., 1995; Kasper, Liu, et al., 2014; Samelson et al., 2016; Touchette et al., 1986) *vs.* the negative Lon (-Lon) selection coefficient, positive Lon (+Lon) selection coefficient, and difference between positive and negative Lon ((+Lon) - (-Lon)) selection coefficients(Thompson et al., 2020). Red dashed lines indicate linear fits to plotted data with corresponding $R^2$ values.

**Supplementary Table 1**. Constructs used in study and relevant references

| | Construct | Reference |
|---|---|---|
| **1** | WTeDHFR | (Garvey & Matthews, 1989; Iwakura et al., 1995; Touchette et al., 1986) |
| **2** | V75R | (Garvey & Matthews, 1989; Samelson et al., 2016) |
| **3** | L8A | (Kasper, Andrews, et al., 2014) |
| **4** | T35D | (Leontiev et al., 1993) |
| **5** | N37S | (Leontiev et al., 1993; Murzina & Gudkov, 1990) |
| **6** | R57H | (Leontiev et al., 1993) |
| **7** | I61A | (Kasper, Andrews, et al., 2014) |
| **8** | I91A | (Kasper, Andrews, et al., 2014) |
| **9** | L110V | (Kasper, Andrews, et al., 2014) |
| **10** | L112V | (Kasper, Andrews, et al., 2014) |
| **11** | V136A | (Kasper, Andrews, et al., 2014) |
| **12** | L156A | (Kasper, Andrews, et al., 2014) |
| **13** | C85A | (Iwakura et al., 1995) |
| **14** | C85S | (Iwakura et al., 1995) |
| **15** | C152E | (Iwakura et al., 1995) |
| **16** | C152S | (Iwakura et al., 1995) |
| **17** | C85A/C152E | (Iwakura et al., 1995) |
| **18** | C85S/C152S | (Iwakura et al., 1995) |
| **19** | AGLA | (Kasper, Andrews, et al., 2014), control |
| **20** | DGLP | control |
| **21** | hDHFR | (Reeve et al., 2019) |
| **22** | P21L | (Cammarata et al., 2017; Tamer et al., 2019) |
| **23** | P21Q | (Watson et al., 2007) |
| **24** | F153L | (Toprak et al., 2012) |
| **25** | F153V | (Toprak et al., 2012) |
| **26** | F153S | (Watson et al., 2007) |
| **27** | A26T | (Cammarata et al., 2017; Tamer et al., 2019; Watson et al., 2007) |
| **28** | A26V | (Toprak et al., 2012) |
| **29** | A26S | (Toprak et al., 2012) |
| **30** | W30R | (Cammarata et al., 2017; Tamer et al., 2019; Watson et al., 2007) |
| **31** | W30G | (Tamer et al., 2019) |
| **32** | M20V | (Watson et al., 2007) |
| **33** | M20I | (Tamer et al., 2019) |

**Supplementary Table 2**. Measured rate constants from off-chip proteolysis with either gel electrophoresis or activity-based readouts.

| Construct | $k_{unfold}$ (from gels) ($s^{-1}$) | $k_{unfold}$ (from activity assays) ($s^{-1}$) |
|---|---:|---:|
| AGLA | 0.0038 | |
| DGLP | <4.4E-05 | |
| L8A | 0.0020 | 0.0046 |
| V75R | 0.00032 | 0.00042 |
| W30R | 0.0030 | |
| WT | 0.00045 | 0.00026 |
| hDHFR | 0.0014 | 0.0012 |

**Supplementary Table 3**. Median SPARKfold-measured rate constants and calculated significance for each construct.

| **Construct** | $\boldsymbol{k}_{\textbf{unfold}}$ **(s$^{-1}$)** | $\boldsymbol{\sigma_k}$ **(s$^{-1}$)** | $\boldsymbol{ln(k_{\textbf{unfold}})}$ **(s$^{-1}$)** | $\boldsymbol{ln(\sigma_k)}$ **(s$^{-1}$)** | $p$ value | significant? |
|---|---|---|---|---|---|---|
| DGLP | 9.1E-05 | 1.1E-05 | -9.3 | 0.1 | 0 | yes |
| AGLA | 0.0047 | 0.0017 | -5.4 | 0.4 | 0 | yes |
| WT | 0.00039 | 0.00026 | -7.8 | 0.7 | 0.84 | no |
| L8A | 0.0031 | 0.0011 | -5.8 | 0.3 | 0 | yes |
| M20I | 0.00041 | 0.00024 | -7.8 | 0.6 | 0.66 | no |
| M20V | 0.00035 | 0.00010 | -7.9 | 0.3 | 0.42 | no |
| P21L | 0.00072 | 0.00029 | -7.2 | 0.4 | 0.0024 | no |
| P21Q | 0.0015 | 0.00089 | -6.5 | 0.6 | 0.0001 | yes |
| A26S | 0.0016 | 0.00057 | -6.4 | 0.3 | 0 | yes |
| A26T | 0.0033 | 0.00083 | -5.7 | 0.3 | 0 | yes |
| A26V | 0.0020 | 0.00077 | -6.2 | 0.4 | 0 | yes |
| W30G | 0.0010 | 0.00068 | -6.9 | 0.7 | 0.0082 | no |
| W30R | 0.0021 | 0.00076 | -6.2 | 0.4 | 0 | yes |
| T35D | 0.0011 | 0.00070 | -6.8 | 0.6 | 0.0016 | no |
| N37S | 0.0014 | 0.00071 | -6.6 | 0.5 | 0.0027 | no |
| R57H | 0.00067 | 0.00022 | -7.3 | 0.3 | 0.0002 | yes |
| I61A | 0.0014 | 0.00092 | -6.6 | 0.7 | 0.0003 | yes |
| C85A | 0.00038 | 0.00018 | -7.9 | 0.5 | 0.91 | no |
| C85A_C152E | 0.0014 | 0.00078 | -6.6 | 0.6 | 0.0015 | yes |
| C85S | 0.00051 | 0.00030 | -7.6 | 0.6 | 0.20 | no |
| C85S_C152S | 0.00053 | 0.00021 | -7.5 | 0.4 | 0.078 | no |
| I91A | 0.00052 | 0.00031 | -7.6 | 0.6 | 0.18 | no |
| L112V | 0.0023 | 0.0014 | -6.1 | 0.6 | 0.0002 | yes |
| V136A | 0.0030 | 0.0019 | -5.8 | 0.6 | 0.0004 | yes |
| C152E | 0.0011 | 0.00026 | -6.8 | 0.2 | 0 | yes |
| C152S | 0.00050 | 0.00027 | -7.6 | 0.5 | 0.099 | no |
| F153L | 0.0031 | 0.0012 | -5.8 | 0.4 | 0.0003 | yes |
| F153S | 0.0026 | 0.00059 | -6.0 | 0.2 | 0 | yes |
| F153V | 0.0046 | 0.0012 | -5.4 | 0.3 | 0.0002 | yes |
| L156A | 0.0056 | 0.0041 | -5.2 | 0.7 | 0.0003 | yes |

**Supplementary Table 3**. Median SPARKfold-measured rate constants and calculated significance for each construct.

| Construct | $\Delta G_{unf}$ (kcal/mol) | $\Phi_c$ | Reference |
|---|---|---|---|
| WT | 6.6±0.3 | – | (Kasper, Liu, et al., 2014) |
| L8A | 4.5±0.1 | 0.05±0.1 | (Kasper, Liu, et al., 2014) |
| I61A | 5.0±0.4 | 0.69±0.09 | (Kasper, Liu, et al., 2014) |
| C85S | 5.4±0.6 | | (Iwakura et al., 1995) |
| I91A | 6.0±0.2 | 0.5±0.2 | (Kasper, Liu, et al., 2014) |
| L112V | 4.9±0.2 | -0.3±0.3 | (Kasper, Liu, et al., 2014) |
| V136A | 5.7±0.1 | 0.5±0.2 | (Kasper, Liu, et al., 2014) |
| C152E | 5.7±0.4 | | (Iwakura et al., 1995) |
| L156A | 3.7±0.1 | 0.24±0.07 | (Kasper, Liu, et al., 2014) |
| C85A/C152E | 5.0±0.4 | | (Iwakura et al., 1995) |
| C85S/C152S | 4.9±0.4 | | (Iwakura et al., 1995) |
